# Lactate and pyruvate promote cellular stress resistance and longevity through ROS signaling

**DOI:** 10.1101/542316

**Authors:** Arnaud Tauffenberger, Hubert Fiumelli, Salam Almustafa, Pierre J. Magistretti

## Abstract

L-lactate, for long considered a glycolytic end-product, is now recognized as an important energy substrate. Moreover, it appears that its role is not limited to energy production but also as a signal for neuroprotection and synaptic plasticity. Using a model of neuroblastoma cells and the nematode *C. elegans* we investigated the cellular mechanisms underlying this protective role of L-lactate. We found that L-lactate promotes a mild Reactive Oxygen Species (ROS) induction that translates into activation of antioxidant defenses and pro-survival pathways such as PI3K/AKT and Endoplasmic Reticulum (ER) chaperones. This hormetic mechanism provides protection against oxidative stress in both cells and nematodes. Furthermore, a mild ROS induction by lactate also promotes longevity in *C. elegans*.

## Introduction

Neurons integrate many signals that translate into high energy needs, leading to the use of approximately 80-90% of the total energy consumed by the brain^1^. Neurons were initially believed to rely exclusively on glucose taken up directly from the circulation, but it is now clear that glial cells, and in particular astrocytes, have a critical role in supporting the metabolic needs of neurons, providing trophic signals and energy metabolites^2,3^. Astrocytes are the main cellular uptake site of blood-borne glucose, which they process mainly through aerobic glycolysis to produce L-lactate (herein referred to as lactate). Neurons, in contrast, rely mainly on oxidative metabolism and import astrocyte-derived lactate to support their high energetic demand in a metabolic coupling mechanism known as astrocyte-neuron lactate shuttle (ANLS)^4^. In neurons, lactate is converted to pyruvate which is further oxidized in the mitochondria by the TCA cycle leading to ATP production^5^. In addition to its role in energy metabolism, lactate also serves as a signal in a variety of processes such as synaptic plasticity and memory consolidation^6,7^. It is also well established that lactate is protective against different insults including glutamate excitotoxicity and ischemia-reperfusion^8,9^.

Mitochondria tightly regulate cellular processes involved in energy production and homeostasis. These organelles use O_2_ to oxidize NADH/FADH_2_ produced by the TCA cycle to generate a series of electron transfers through protein complexes resulting in the establishment of an electrochemical gradient that powers the production of ATP. Reactive Oxygen Species (ROS) are physiological by-products of normal cellular respiration, but their levels can dramatically increase when the respiratory chain is dysfunctional. Thus, mitochondrial dysfunctions during aging or cellular stress are thought to be involved in neurodegenerative diseases, including Alzheimer’s and Parkinson’s disease^10^. The energetic deficits during aging are also correlated with the loss of proteostasis, a set of processes controlling the homeostasis of the biogenesis, folding, trafficking, and degradation of proteins^11,12^.

Until recently ROS were considered a hallmark of oxidative stress leading to cellular dysfunction and neurodegeneration^13,14^. Growing evidence now indicates that ROS may also act as signaling molecules in physiological processes. Indeed, exposure for short time periods or low concentrations of ROS contributes to increased lifespan in multiple organisms^15-17^ in a pro-survival mechanism called hormesis or mito-hormesis^18-20^.

Here we show that lactate promotes resistance to oxidative stress in both mammalian cells and *C. elegans*, and increases the longevity of *C.elegans,* where pyruvate reproduces the protective effects mediated by lactate. We found that lactate supplementation induces a moderate elevation in ROS levels and transcription of genes belonging to pro-survival pathways, including the IGF-AKT/PI3K and the endoplasmic reticulum stress pathways. These observations suggest that lactate, and to a lesser extent pyruvate, supports resistance to cellular stress and promotes longevity through a mild hormetic increase in oxidative stress.

## Results

### Lactate pre-treatment reduces cell death induced by oxidative stress

Lactate has been shown to play a role in enhancing neuronal survival^9,21^. These studies indicated that lactate promotes an increased ATP production and a better Ca^2+^ buffering in a model of excitotoxicity. We, therefore, set out to investigate whether lactate could also promote cell protection using SH-SY5Y neuroblastoma cells, a widely used model of cell toxicity and neurodegeneration^22^. This cell line has also been used to investigate the regulation of gene expression by lactate^23^. We treated the cells with a high concentration of hydrogen peroxide (H_2_O_2_), one of the primary cellular ROS, and assessed cell death using the Trypan Blue exclusion method **(Fig. 1a)**. Initial results showed that a concentration of 150 µM of H_2_O_2_ resulted in 70-80 % of cell death after 24h of oxidative stress. Under these conditions, we tested whether supplementing the medium with lactate (20 mM) could promote cell survival. Co-application of lactate together with H_2_O_2_ did not counteract the toxic effects of H_2_O_2_-induced oxidative stress, while pyruvate co-treatment massively reduced cell death. The ROS scavenging effect of pyruvate, results from direct chelation of H_2_O_2_ leading to its inactivation in the cell culture media^24,25^, and is therefore artefactual in terms of cellular processes **(Fig. 1b).**

**Figure 1:**
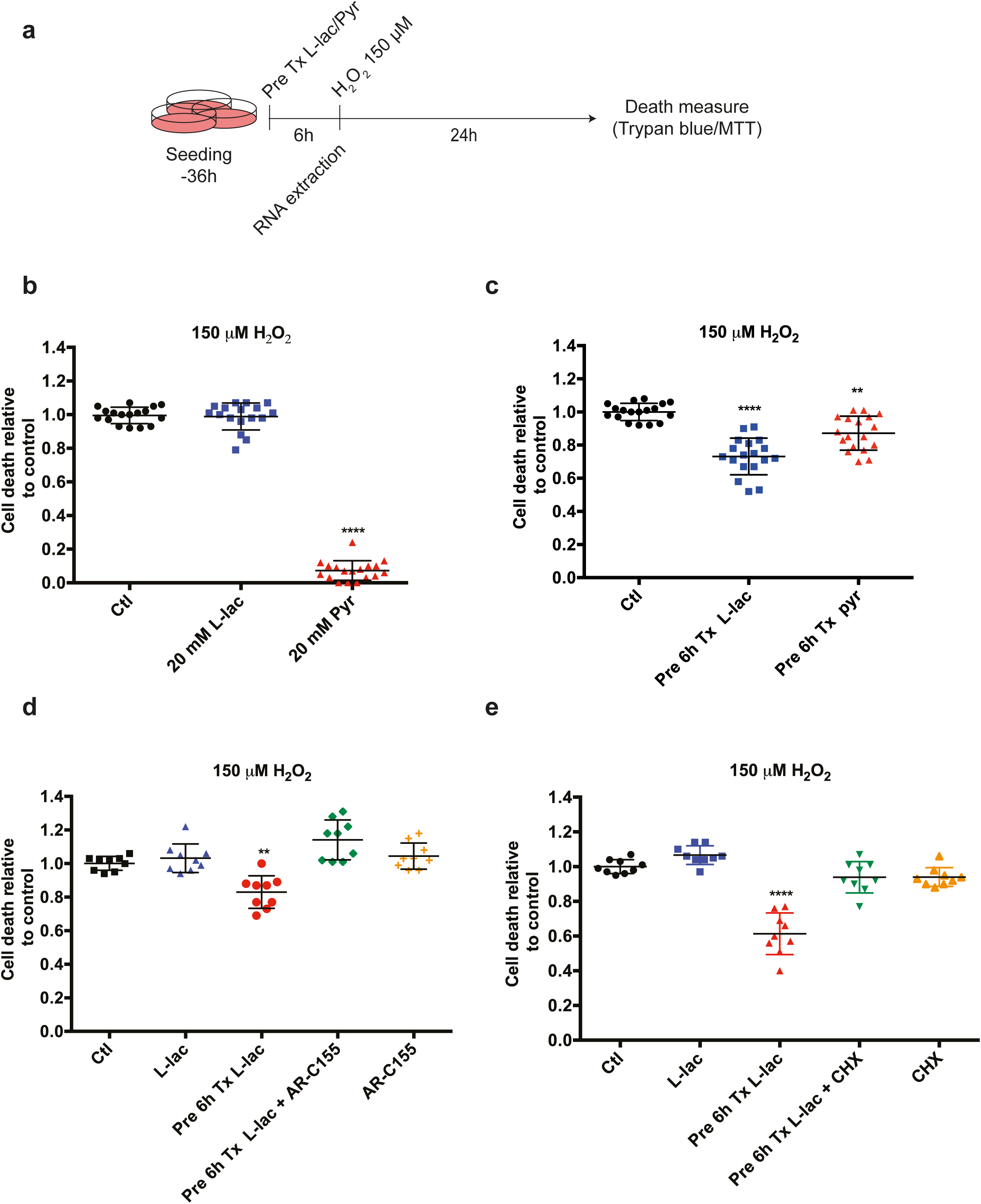
Pre-treatment with lactate increases resistance to oxidative stress. **a)** Schematic representation of the experimental procedures used in this study with the SH-SY5Y cells. **b)** Measures of cell death after co-application of 150 µM H_2_O_2_ with lactate or pyruvate. **c)** Measures of cell death after oxidative stress induced by of 150 µM H_2_O_2_ following a 6h pre-treatment with lactate or pyruvate. **d-e)** Measures of cell death after oxidative stress induced by 150 µM H_2_O_2_ upon pre-treatment with lactate or pyruvate and application of the MCT1 inhibitor AR-C155858 (D) or 1 µM cycloheximide (E). Each data point represents one measurement. Bars are the average ± SEM. All calculations for statistical significance were completed using a non-parametric, one-way ANOVA with multiple comparisons, n= 12-15 from 4-5 independent experiments. **p<0.01, ****p<0.0001

We next tested a possible effect of lactate and pyruvate as pre-treatment on cell death triggered by H_2_O_2_ exposure. Lactate media supplementation for 6h prior to oxidative stress exposure resulted in significantly increased survival (26,8%) **(Fig. 1c and Supplementary Fig. 1a)** while limited or no effect was observed with pyruvate. These observations were not dependent on the detection method for cell death, as MTT assay offered comparable results to Trypan Blue exclusion **(Supplementary Fig. 1b).** An osmolarity effect of 20 mM lactate can be excluded as pre-treatment with 20 mM NaCl, D-glucose or Na-gluconate did not affect cell survival upon H_2_O_2_ treatment **(Supplementary Fig. 1c)**. Finally, lactate was also able to rescue cell death upon treatment with two other oxidative stress inducers, tert-butyl hydroperoxide and sodium arsenite **(Supplementary Fig. 1d, e**), suggesting that the observed protective effect is not limited against the stress triggered by H_2_O_2_.

Lactate and pyruvate are transported through cells via a family of plasma membrane monocarboxylate transporters (MCT). To determine whether cellular uptake of lactate was required for protection against oxidative stress, we examined the lactate-dependent decrease in H_2_O_2_-induced cell death in the presence of the MCT blocker AR-C155858. Inhibition of MCT was found to decrease the protective effect of lactate **(Fig 1d)** on H_2_O_2_-evoked oxidative stress. Similar attenuation of lactate’s effect was also observed using UK5099, another MCT inhibitor **(Supplementary Fig. 1f)**. These data indicate that the entry of lactate into cells is necessary for its protective effects on oxidative stress-induced cell death.

To establish whether lactate enhanced cell survival through *de novo* protein synthesis, we investigated the protective effects mediated by lactate in the presence of the translation blocker cycloheximide. Interfering with the translation machinery prevented the cell survival effect of lactate on oxidative stress-evoked cell death **(Fig. 1e)**.

### Lactate reduces cell death through induction of pro-survival and proteostatic pathways

To further characterize the molecular underpinnings of the protective effect of lactate, we performed a differential gene expression analysis on SH-SY5Y cells. Using whole transcriptome RNA sequencing (RNAseq), resulting in the identification of the expression of over 13000 genes, we found that treatment of neuroblastoma cells with lactate or pyruvate respectively affected the expression of 1261 (640 up-regulated and 621 down-regulated) or 1916 genes (958 up-regulated and 958 down-regulated). The comparison between lactate- and pyruvate-modulated genes suggests that both metabolites affect a large set of transcriptional changes in a similar way (1281 similarly regulated genes as compared to mock treatment). This comparison also enabled us to identify specific transcriptional responses that are unique to lactate **(Fig. 2a, b)**. To have a better grasp of the high-level functions and mechanisms regulated by lactate, we identified over-represented biological processes as well as molecular functions and cellular components from the list of genes specifically regulated by lactate using gene and pathway ontology enrichment analyses (see materials and methods). The most significantly enriched gene ontology terms for differentially expressed genes (DEG) evoked by lactate included cell metabolism, growth, and survival while over-representation analysis of KEGG pathways revealed the involvement of PI3K, mTOR signaling and protein processing in ER pathways **(Fig. 2c)**. These observations are consistent with previous findings suggesting that lactate-mediated neuroprotection involves the PI3K signaling pathway^9^. mTOR signaling is a major pathway involved in cell growth and survival in different cell types^26,27^. Among the genes specifically affected by lactate (as opposed to pyruvate) pre-treatment and included in one of the three enriched pathways, we identified TSC2 and S6K (mTOR signaling), AKT1 (PI3K pathway), as well as ATF4, TRAF2, GRP78/BiP and XBP1 (ER processing) **(Supplementary Fig. 2a-g)**. GRP78/BiP and XBP1 are part of the unfolded protein response in the ER (UPR^ER^) and promote protein homeostasis^28,29^. Additionally, we identified several ER chaperones uniquely regulated by lactate, including DNAJA2, DNAJC5, and DNAJC10 **(Supplementary Fig. 2h-k)**. To verify the involvement of these important pathways in the protective effects induced by lactate, we first examined the role of the PI3K/Akt pathway by treating the cells with a potent PI3K inhibitor. Co-application of lactate and LY294002 blocked the rescue by lactate of oxidative stress-evoked cell death **(Fig. 3a).** The contribution of the UPR^ER^ in the lactate-dependent protection against ROS toxicity was examined using quercetin, a flavonol known to reduce UPR^ER^ activity^30^. Quercetin decreased the protective effect by lactate after H_2_O_2_ treatment **(Fig. 3b)**. Taken together, these results establish that lactate promotes cell survival against oxidative stress through the induction of key homeostatic pathways that include PI3K, mTOR and ER protein processing.

**Figure 2:**
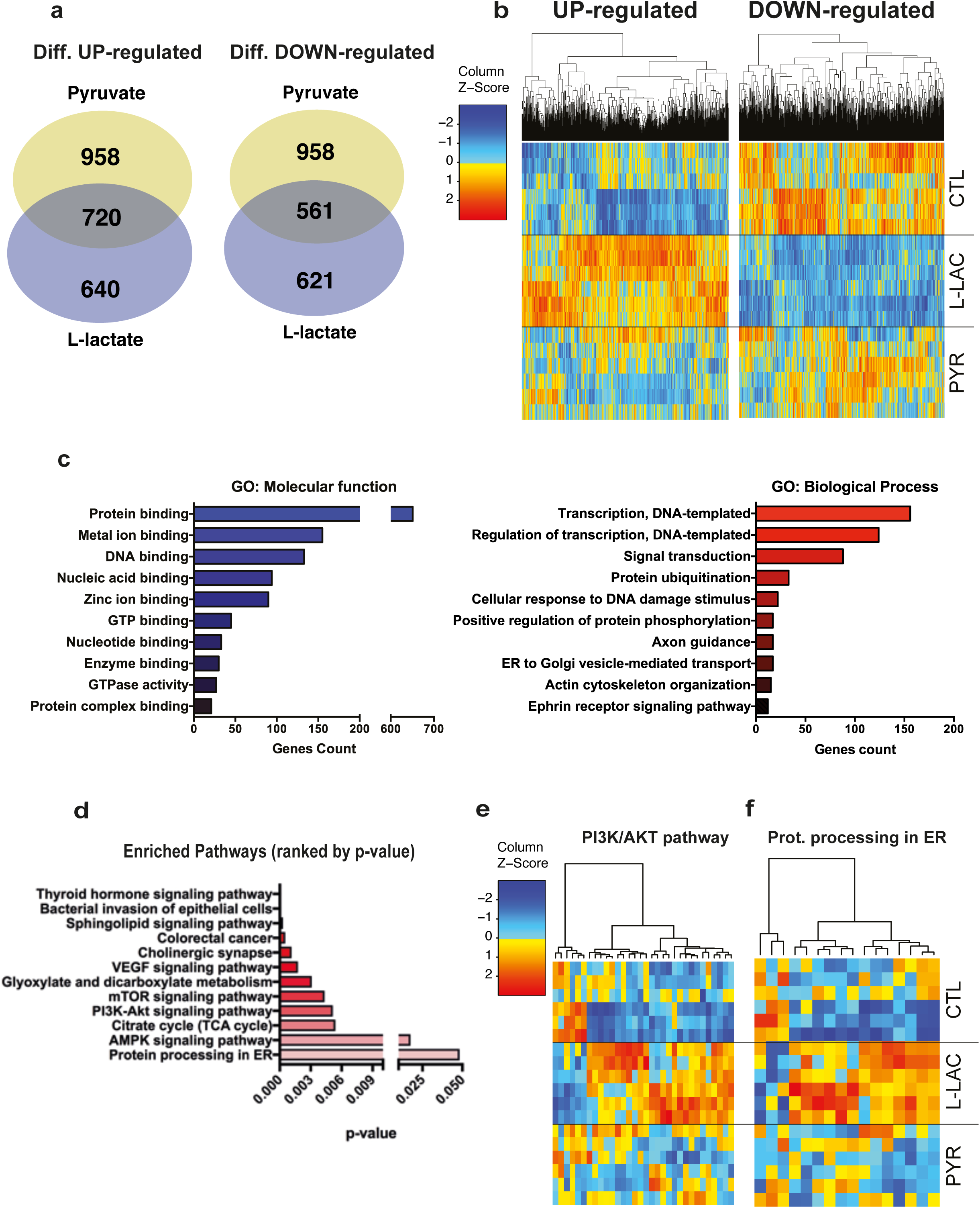
RNA sequencing of SH-SY5Y cells revealed enrichment in key growth-survival pathways. **a)** Venn diagram showing the overlap between the numbers of significant differentially up- and down-regulated genes compared to control by lactate and pyruvate. SH-SY5Y cells were treated for 6h with 20mM lactate or pyruvate. Cut-off for significance was p<0.01. **b)** Heat map clustering of genes differentially regulated by lactate, compared to control and pyruvate treated cells. The map represents expression levels of each transcript (count per million), and z-score represents normalized expression of each transcript across the different conditions. **c-d)** Molecular functions and biological processes GO terms **c)** and KEGG pathways **d)** over-representation analyses. GO terms are ranked by the number of genes while enriched pathways are ranked by increasing p-value (Fisher’s test). **e-f)** Heatmap clustering for KEGG-PI3K/AKT **e)** and KEGG-Protein processing in ER **f)**. Each map represents the list of genes specifically regulated by lactate and associated with PI3K/AKT or Protein processing in the ER. The map represents expression levels of each transcript (count per million), and z-score represents normalized expression of each transcript across the different conditions.

**Figure 3:**
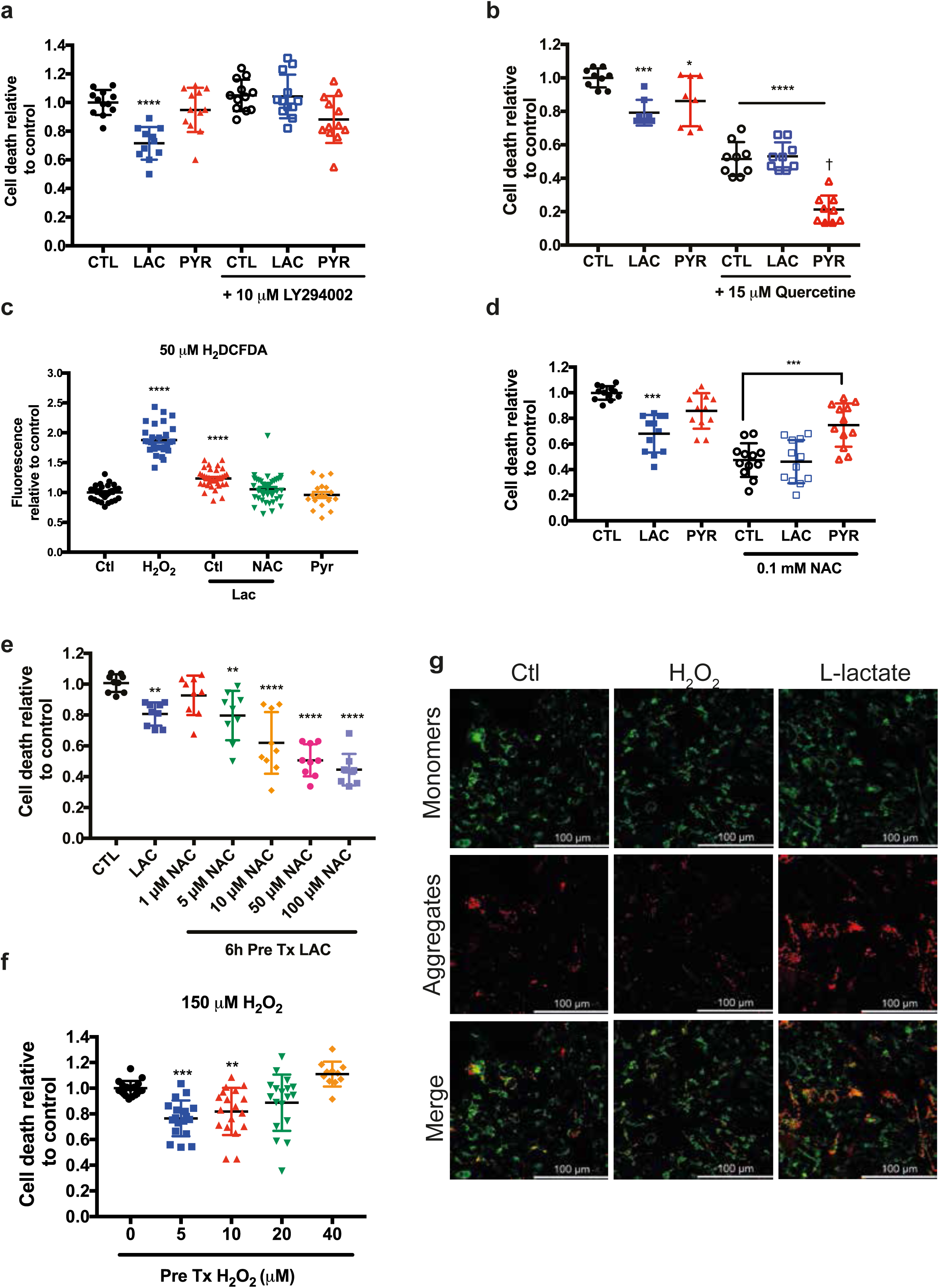
PI3K and UPR^ER^ pathways mediate lactate mediated cell survival, through ROS production. **a)** Relative cell death assessed by trypan blue exclusion following oxidative stress induced by 150 µM H_2_O_2._ Prior to oxidative stress exposure, SH-SY5Y cells were co-treated for 6h with 10 µM LY2940002 and lactate or pyruvate. **b)** The measure of cell death after 150 µM H_2_O_2_ treatment upon pre-treatment with lactate and pyruvate. Cells were co-treated with UPR^ER^ inhibitor (Quercetine – 15µM) with 20 mM lactate or pyruvate. **c)** The measure of ROS level using H_2_DCFDA upon 150 µM H_2_O_2_, 20mM lactate or pyruvate and 50 µM N-acetylcysteine. **d)** Relative cell death measurements following oxidative stress induced by 150 µM H_2_O_2._ Prior to oxidative stress exposure, cells were co-treated for 6h with 100 µM N-acetyl-cysteine lactate or pyruvate. **e)**. Relative cell death measurements following oxidative stress induced by 150 µM H_2_O_2._ Prior to oxidative stress exposure, cells were co-treated for 6h with increasing doses of N-acetyl-cysteine and 20mM lactate. **f)** Measure of cell death after 150 µM H_2_O_2_ and pre-treated with low doses of H_2_O_2_. **g)** Confocal images of SH-SY5Y cells stained with JC1 (2.5 µM). Cells were treated with 150 µM H_2_O_2_ for 30 min or with 20mM lactate for 6h. Each data point represents one measurement. Bars are the average ± SEM. All calculations for statistical significance were completed using a non-parametric, one-way ANOVA with multiple comparisons, n= 12-15 from 4-5 independent experiments. *p<0.05, **p<0.01, ***p<0.001, ****p<0.0001, ^†^(p<0.0001 compared to Quercetine treated ctl)

### Lactate reduces cell death through ROS signaling

It is worth noting that most of these pathways have been linked to ROS signaling. Detailed examination of the transcriptomics data, focusing on ROS detoxification enzymes, provided evidence that the expression of HIFα transcript is increased by lactate **(Supplementary Fig. 2l)** while that of NRF2 was increased by both lactate and pyruvate pre-treatment **(Supplementary Fig. 2m)**. The effectiveness of a pre-treatment as a potential protective mechanism suggests the induction of a protective response or hormesis that would reduce the impact of lethal stress later on^18,19,31^. Work by others has shown that lactate promotes ROS induction^32,33^, suggesting that this mechanism could mediate the observations reported here.

To study the involvement of ROS in the protective effect of lactate, we measured ROS production in cells pre-treated with lactate or pyruvate using 2’,7’-dichlorodihydrofluorescein di-acetate (H_2_DCFDA), a fluorescent dye detecting reactive oxygen intermediates. A significant increase in ROS production was observed 6h after lactate treatment when compared to both control and pyruvate treated cells and this increase was lost upon treatment with antioxidant N-acetyl-L-cysteine (NAC, 0.1mM) **(Fig. 3c and Supplementary Fig. 1g)**^34^. In the presence of NAC, lactate was not able to reduce cell death further indicating that cell protection evoked by lactate requires an increase in ROS levels **(Fig. 3d)**. Of note is the fact that NAC decreased by itself the overall cell death upon H_2_O_2_ treatment. We confirmed our hypothesis by adding increasing concentrations of NAC to cells pre-treated with lactate. A low concentration of NAC (1 µM), which was not able on its own to reduce overall cell death by H_2_O_2_, nevertheless blocked the protective effect of lactate **(Fig. 3e)**. These data confirm that ROS induction by lactate is required for cell survival upon oxidative stress. Furthermore, we were able to mimic the lactate-mediated protection against oxidative stress using pre-treatment with low doses of H_2_O_2_ **(Fig. 3f)**.

Given the role of the mitochondria in ROS production, we sought to examine if lactate could affect mitochondrial function. Using staining with JC1, a fluorescent mitochondria indicator forming red shifted J-aggregates at high membrane potentials^35^, we observed an increase in mitochondrial potential (red aggregates) upon lactate treatment **(Fig. 3g)**.

In contrast, high doses of H_2_O_2_ dramatically reduced mitochondrial membrane potential (green staining). These results indicate that lactate increases mitochondrial respiration resulting in a mild induction of ROS levels, while a high H_2_O_2_ concentration disrupts membrane potential, causing mitochondrial dysfunction and cell death. Besides the regulation of transcription factors NRF2 and HIF1α expression, the RNAseq analysis did not reveal changes in the expression of effector enzymes controlling ROS detoxification after 6h. However, a measure of the expression of SOD1, PRDX5, GSTM4 and GPX3 3h after the end of exposure to lactate or pyruvate showed a significant increase. **(Supplementary Fig. 2n-q).** Altogether, the effects reported here strongly suggest that lactate supplementation induces a mild ROS production, which in turn activates pro-survival pathways including mTOR, PI3K and ER protein processing.

### Lactate and pyruvate delay aging-evoked phenotypes in *C. elegans*

We next took advantage of *C. elegans’s* simple genetics and short lifespan to investigate, *in vivo*, the role of lactate in stress resistance and longevity in relation to ROS signaling. First, in order to investigate the effect of lactate on nematode physiology, we supplemented their bacterial diet with lactate or pyruvate while inducing oxidative stress by exposure to the natural compound juglone^36^. Juglone is a natural compound from the black walnut tree that increases intracellular concentrations of superoxide

After 8 hrs of exposure to oxidative stress, animals supplemented in their diet with high concentrations of lactate or pyruvate (100 mM) had higher survival rates than control nematodes **(Fig. 4a)**. It is worth noting that the concentration of metabolites in the media is not the one found in the nematode, as shown in different studies investigating how the energy metabolism influences longevity^37,38^. Lower concentrations of lactate or pyruvate (50 and 10 mM) did not increase survival, suggesting that the protective mechanisms underlying the stress resistance were not triggered **(Supplementary Fig. 3a, b)**. “Beneficial” levels of stress, or hormesis, can prime organisms and increase their resilience, sometimes resulting in increased life span^31^. Therefore, we also measured the longevity of *C.elegans* in the presence of monocarboxylates. Interestingly, a low concentration (10 mM) of lactate or pyruvate increased the lifespan of wild-type animals **(Fig. 4b)** whereas higher concentrations (50 mM and 100 mM) reduced lifespan **(Supplementary Fig. 3c, d)**. The increased stress resistance and longevity are not caused by secondary metabolism of lactate or pyruvate by the nematodes’ bacterial food source; animals grown on heat-killed bacteria with no metabolic activity showed the same response to lactate and pyruvate **(Supplementary Fig. 3a, b and e)**.

**Figure 4:**
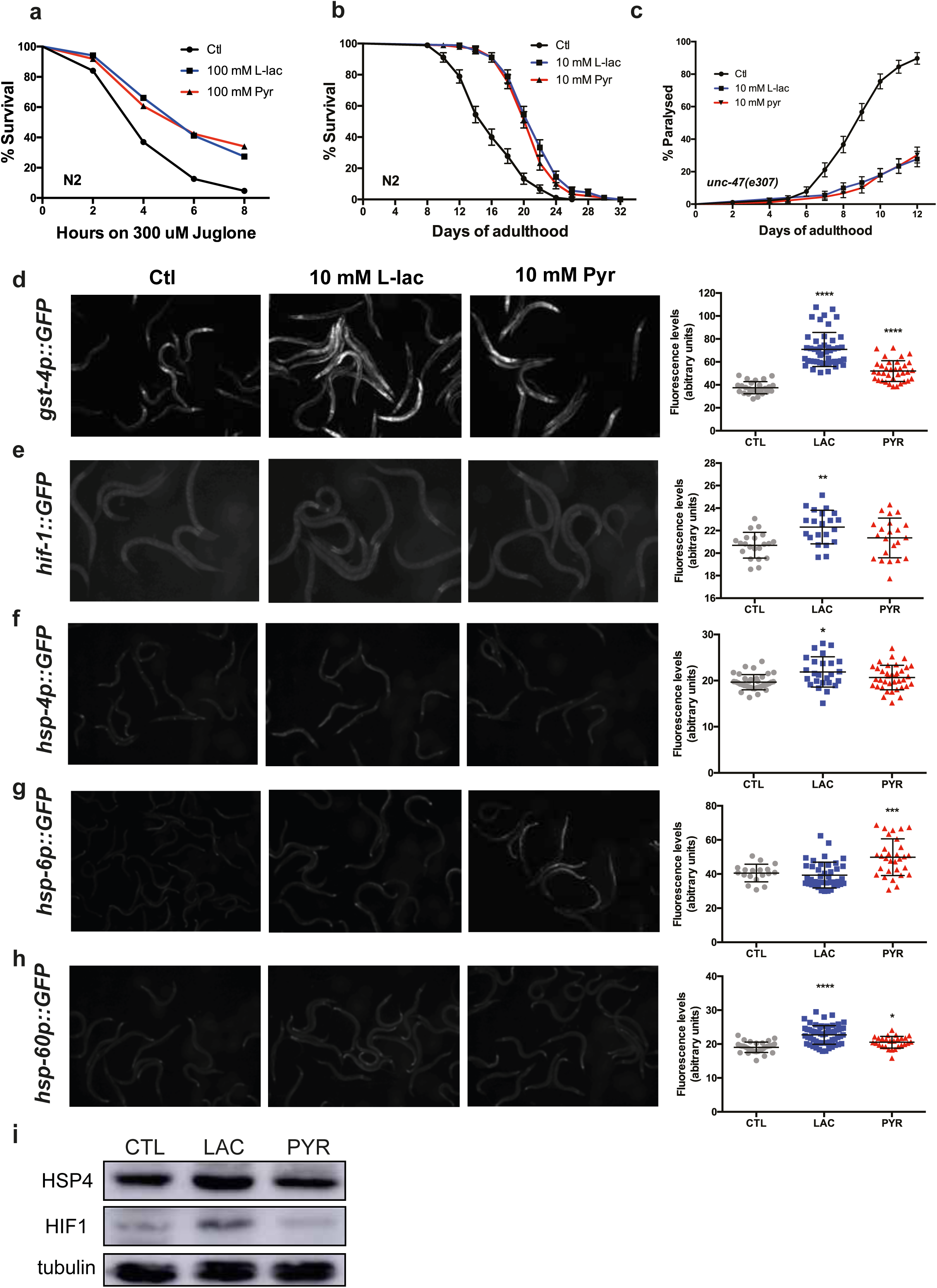
lactate and pyruvate increase oxidative stress resistance and longevity through ROS detoxification and UPR^ER^ activation. **a)** Survival curves of wild-type nematodes supplemented with lactate or pyruvate treated with 300µM juglone **(p<0.0001)** **b)** Lifespan curves of wild-type animals treated with 10 mM lactate or pyruvate. **c)** Paralysis curve of *unc-47(e307)* animals treated with 10 mM lactate or pyruvate. **d-h)** Representative GFP fluorescence images and quantitative measurements of GFP fluorescence intensities in *C.elegans* expressing transcriptional reporters, as indicated in the left margin, treated with 10mM lactate or pyruvate. **i)** Immunoblot analysis of HIF1α and HSP4/GRP78 protein expression in wild-type nematode. Tubulin is used as a loading control. For survival curves p-value < 0.0001 was determined using the Log-Rank (Mantel-Cox) test. 90-100 animals were used per condition for each trial. Experiments were repeated at least three times. See supplementary table I for lifespan statistics. For the quantitative GFP summaries, each data point represents the mean fluorescence value of one animal. Horizontal bars are the average ± SEM. Statistical significance (*p<0.05, **p<0.01, ***p<0.001 and ****p<0.0001) was calculated using a non-parametric, one-way ANOVA with multiple comparison test, n = 30-76 from 3 independent experiments.

One hallmark of aging is an increased prevalence of neuronal dysfunction and we reasoned that increased longevity might be linked to neuroprotective effects of lactate and pyruvate. To test the effect of lactate and pyruvate on age-related neuronal dysfunction, we used *unc-47(e307)* mutant animals which have impaired GABA neurotransmission and develop age-dependent paralysis as a consequence^39^. *unc-47* mutant animals exhibited delayed paralysis onset as well as a slower paralysis progression when supplemented with 10 mM lactate or pyruvate compared to mutant animals grown on control media **(Fig. 4c)**. In a second model, neural communication between neurons and muscles can be tested at the neuromuscular junction (NMJ) with the drug aldicarb, an acetylcholinesterase inhibitor which causes acetylcholine accumulation and paralysis^40^. In the presence of aldicarb, young adults (1 day) are rapidly paralyzed **(Supplementary Fig. 3f)** whereas paralysis at older ages (5 and 9 days) is reduced due to an age-dependent reduction in acetylcholine release. **(Supplementary Fig. 3g, h).** Lactate or pyruvate did not change paralysis in response to aldicarb in young adult day 1 animals **(Fig. S3F)**, but both monocarboxylates showed a higher rate of paralysis in aging *C.elegans* **(Supplementary Fig. 3g, h),** suggesting these animals may be physiologically younger.

In sum, lactate and pyruvate have significant effects on *C. elegans* including protection from oxidative stress, life-span extension, and reduced aging-related neural dysfunction whether caused by mutations or pharmacological treatment.

### Lactate and pyruvate stimulate ROS detection pathways, UPR^ER^ and UPR^mt^ in *C. elegans*

To determine if the pathways identified in the screen performed on mammalian cells are also involved in L-lactate mediated protection in the nematode, we investigated several well-described transcriptional/translational reporters’ induction by lactate and pyruvate. The expression of the reporter glutathione-S-transferase (*gst-4*), a downstream target of the transcription factor NRF2/*skn-*1 known to contribute to cellular homeostasis against ROS and ER stress^41^, was strongly induced in animals supplemented with lactate or pyruvate. Both metabolites also induced the expression of the ER stress chaperone *hsp-4p*::GFP and the hypoxia-inducible factor *hif-1p*::*hif-1*::GFP reporters **(Fig. 4d-f)**. Lactate- or pyruvate-enriched diet was also capable of inducing expression of mitochondrial chaperones *hsp-60* or *hsp-6,* respectively **(Fig. 4g, h)**. However, expression of *daf-16, sod-3, lgg-1* and *crtc-1* reporters was not influenced by lactate or pyruvate supplementation **(Supplementary Fig. 4a-d)**. These results indicate that pathways involved in mammalian cell protection including ROS detoxification mechanisms, HIF1α, and ER stress responses, identified in SH-SY5Y cells, are activated by lactate or pyruvate in the nematode. Additionally, we determined that the mitochondrial unfolded protein response (UPR^mt^) is induced by lactate and pyruvate supplementation in the nematode **(Fig. 4g, h)**.

### Lactate and pyruvate increase oxidative stress resistance through multiple pro-survival pathways

Following on our observations that lactate induces ROS in SH-SY5Y cells and the observation of increased activity of *gst-4* in the worm **(Fig. 4d)**, we assessed the role of ROS production upon lactate and pyruvate in stress response in the nematode. The use of 10 mM NAC on worms supplemented with 100 mM lactate or pyruvate in the diet abolished the resistance to juglone **(Fig. 5a)**. Furthermore, in line with our findings on neuroblastoma cells **(Fig. 3c)**, we found an increase in the H_2_O_2_ levels in nematodes when these animals were supplemented with lactate or pyruvate compared to untreated controls **(Fig. 5b).**

**Figure 5:**
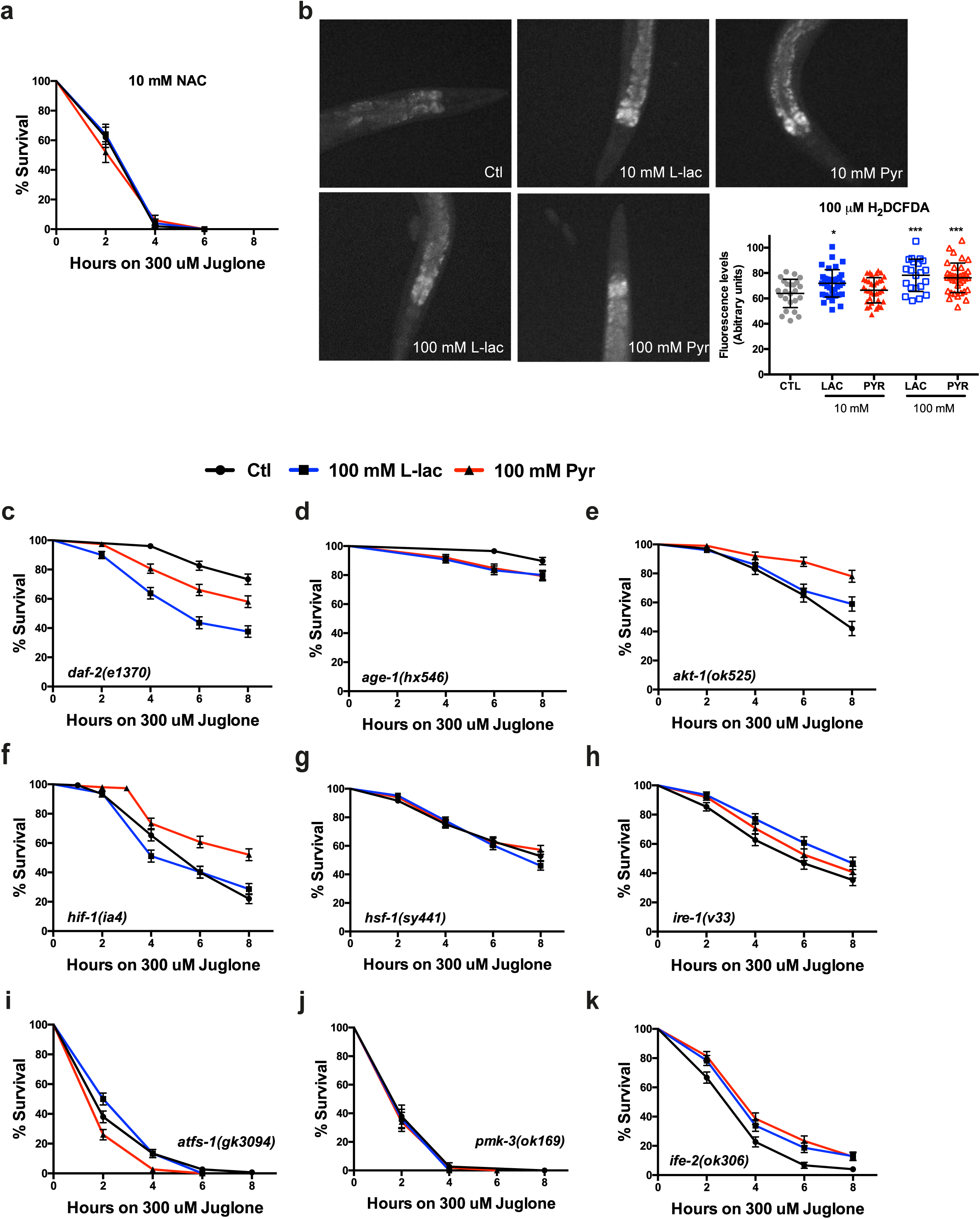
lactate and pyruvate increase oxidative stress resistance through Insulin/IGF pathway and ROS signalling. **a)** Survival curves of wild-type nematodes treated with 300µM juglone and supplemented with 10 mM N-acetyl cysteine (NAC) together with 100mM lactate or pyruvate. **b)** Representative H_2_DCFDA fluorescence images and quantitative measurements of H_2_DCFDA fluorescence intensities in wild-type nematodes supplemented with 10mM or 100mM lactate or pyruvate. **c-k)** Survival curves of mutant nematodes treated 300µM juglone and supplemented with 100mM lactate or pyruvate. Mutated genes are indicated as an inset in the lower left or right corner of each graph. Statistical significance for the survival curves was determined by comparing curves of monocarboxylate-supplemented nematodes to controls using the Log-Rank (Mantel-Cox) test. 90-100 animals were used per condition for each trial. Experiments were repeated at least three times. See supplementary table I for lifespan statistics. For the quantitative fluorescence measurements summary, each data point represents the mean fluorescence value of one animal. Horizontal bars are the average ± SEM. Statistical significance (*p<0.05 and ***p<0.001) was calculated using a non-parametric, one-way ANOVA with multiple comparison test, n = 30-50 from 3 independent experiments.

To understand how ROS induction promotes oxidative stress resistance, we performed a *C. elegans* screen with candidate genes. We chose a broad array of mutant animals with defective metabolism, stress response, and transcriptional regulation pathways. To identify the signaling pathways pyruvate and lactate act through, we exposed the mutant animals to oxidative stress with juglone in the presence of ordinarily protective concentrations of lactate and pyruvate. If the monocarboxylates are protective in the absence of a given signaling pathway, then that pathway is unlikely to be the cause of protection.

The Insulin/IGF signaling (IIS) is a well-conserved longevity pathway that responds to metabolic changes and triggers the activation of pro-longevity mechanisms^42^. Three components of the (IIS) appeared to be involved in the stress resistance mediated by both monocarboxylates: The knockdown of IGF receptor *daf-2* and the kinases *age-1* and *akt-1* limited stress resistance by lactate or pyruvate. Interestingly, *age-1*, the ortholog of PI3K, and *akt-1* were also identified in our screen in SH-SY5Y cells **(Fig. 5c-e).** Surprisingly, the canonical downstream transcription factor of this pathway, *daf-16,* did not influence lactate- or pyruvate-mediated resistance to juglone **(Supplementary Fig. 5a)**.

Hypoxia-inducible factor 1 (*hif-1*) regulates resistance to hypoxia and longevity upon ROS induction^43,44^. Our mutant screen revealed that a knockdown of *hif-1* blocked the protective effect by lactate and reduced the effect of pyruvate upon oxidative stress **(Fig. 5f)**. This is consistent with our observation that *hif-1* is upregulated upon supplementation with lactate **(Fig. 4e).**

We also observed that *hsf-1, ire-1, and atfs-1* mutants, involved in protein homeostasis under various cellular stresses (heat, ROS, protein misfolding), blocked lactate- or pyruvate-evoked stress resistance **(Fig. 5g-i)**. All promote the expression of various chaperones^45^ or unfolded protein response (UPR) activation to promote stress resistance^46,47^. *ire-1* expression was also enriched in our transcriptomic analysis in SH-SY5Y cells, and it has been shown that NRF2 can promote its activation upon oxidative stress^41,48^. However, a mild mitochondrial electrons transport chain (ETC) disruption by complex I and III mutants, *nuo-6* and *isp-1,*^17,49^ did not influence survival increase evoked by lactate and pyruvate under juglone treatment **(Supplementary Fig. 5b, c)** More surprisingly, knockdown of *pmk-3,* a p38 MAPK, and if *ife-2*, a translation initiation factor (eIF-4E) also blocked stress resistance by lactate and pyruvate **(FIG. 5j, k)**. *pmk-3* is known to regulate longevity induced by mitochondrial disruption and axonal regeneration pathways^50,51^, consistent with our overall observations in cells and the ROS induction observed in the nematode. However, knockdown of *pmk-3* canonical upstream control *dlk-1* did not influence lactate- or pyruvate-mediated stress resistance **(Supplementary Fig. 5d)**. *ife-2* is activated by mTOR and promotes new protein translation. The factor loss of function has been linked to increased longevity^52,53^.

Our mutant screen also showed that well-known genes for canonical pathways involved in metabolism, survival and cellular signaling did not influence the protective effects elicited by lactate or pyruvate. **(Supplementary Fig. 5e-o)**.

Overall, the mutant screen supports the idea that lactate and pyruvate mediate stress resistance through HIF1α as well as PI3K/AKT and UPR^ER^ pathways. Furthermore, consistent with our work on SH-SY5Y cells, lactate and pyruvate also stimulate ROS production in the worm.

### Lifespan extension by lactate and pyruvate is dependent on ROS induction

We sought to determine whether the genes involved in lactate- or pyruvate-mediated stress resistance in the worm were also involved in the regulation of the lifespan by both metabolites **(Fig. 6a)**. Consistent with our findings on oxidative stress in SH-SY5Y cells and the nematode, the anti-oxidant NAC prevented the lifespan extension by lactate and pyruvate in wild-type animals **(Fig. 6f)** These observations indicate that lactate- and pyruvate-mediated elevations of ROS levels are required for lifespan extension. We observed that in *hif-1, ife-2* and *pmk-3* mutants prevented the lifespan extension by lactate and pyruvate **(Fig. 6b-d)**. In contrast, most IIS mutants, as well as *hsf-1* and *ire-1* loss of function, did not affect lifespan extension by lactate or pyruvate **(Supplementary Fig. 6a-e)**. Interestingly, lactate and pyruvate reduced the longevity of *daf-2* mutant **(Fig. 6e)**.

**Figure 6:**
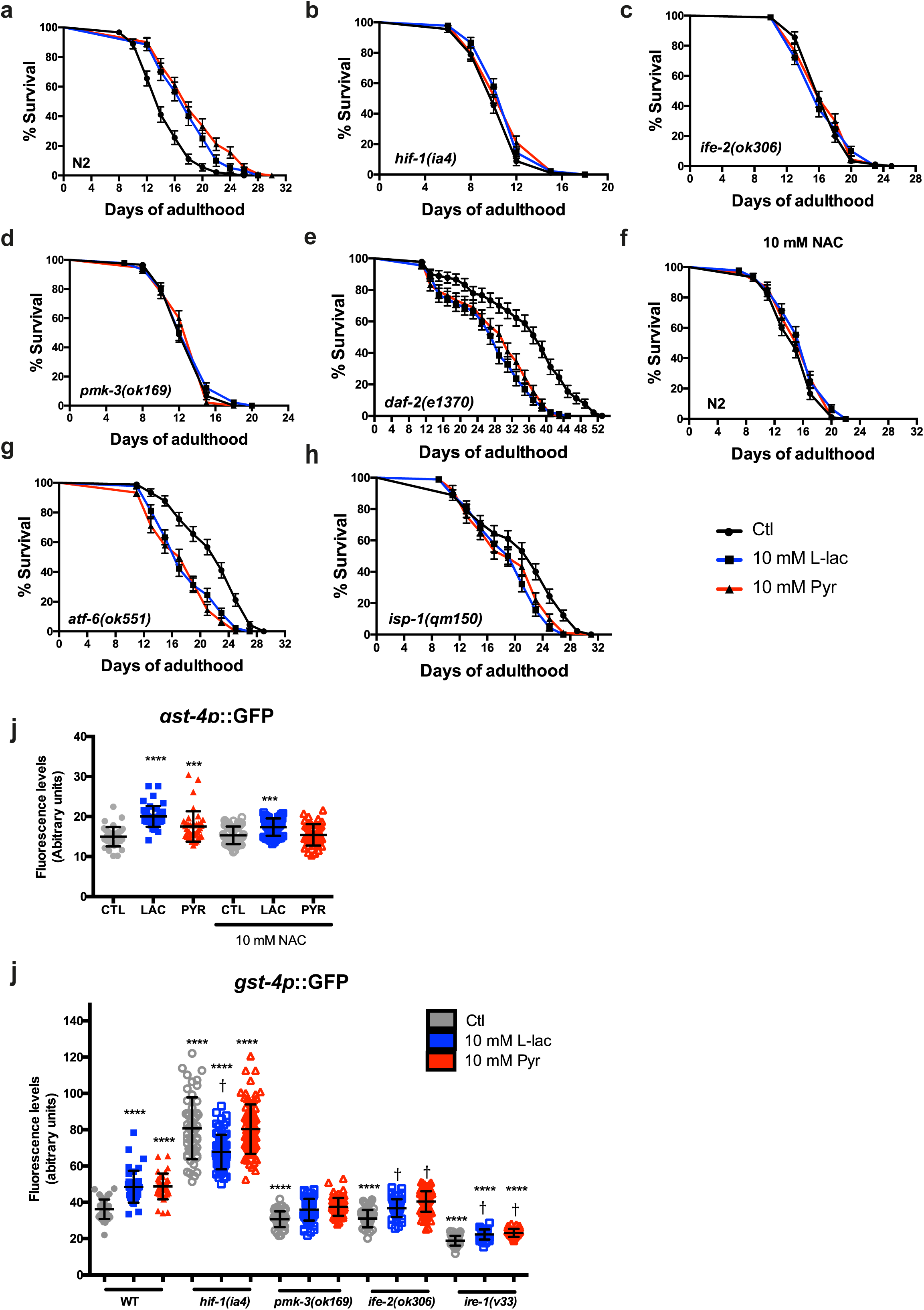
Lifespan extension by lactate and pyruvate depends on mild ROS increases and detoxification mechanisms. **a-h)** Lifespan curves of wild-type (N2) or mutant nematodes treated with 10mM lactate or pyruvate. Wild-type genotype (N2) or mutant genes are indicated as an inset in the lower left corner of each graph. Nematodes were co-treated with 10 mM N-acetyl cysteine (NAC) **f)**. **i-j)** Quantitative summaries of GFP fluorescence in nematodes expressing the transcriptional reporter *gst-4*p::GFP supplemented with 10 mM lactate or pyruvate. **i)** Animals were co-treated with 10mM N-acetylcysteine (NAC) and 10 mM lactate or pyruvate. Statistical significance for the survival curves was determined using the Log-Rank (Mantel-Cox) test. 90 animals were used per condition in each trial. Experiments were repeated at least three times. See supplementary table I for lifespan statistics. For the quantitative fluorescence summaries, each data point represents the mean fluorescence value of one animal. Horizontal bars are the average ± SEM. Statistical significance (****p<0.0001 and ^†^p<0.0001 compared to wild-type and mutant controls, respectively) was calculated using a non-parametric, one-way ANOVA with multiple comparison test, n = 30-70 from 3 independent experiments.

Because results in the nematode show different genetic requirements regarding the genes involved in stress resistance and longevity, we next sought to determine whether other genes from enriched pathways such as UPR^ER^ and UPR^mt^ could participate in the longevity effects by lactate and pyruvate. We observed that the mitochondrial ETC mutant *isp-1* **(Fig. 6g)** and the UPR^ER^ mutant *atf-6* **(Fig. 6h)** were both required for the lifespan extension by lactate and pyruvate, whereas the UPR^mt^ mutant *atfs-1*^54^ **(Supplementary Fig. 6f)** did not influence the longevity increase evoked by lactate and partially reduced that by pyruvate. These data indicate that although ROS signaling similarly mediates stress resistance and longevity by lactate or pyruvate, there are differences in the genetic requirements for each phenotype and the concentration necessary for the protective effect reported here.

Lactate is converted to pyruvate by lactate dehydrogenase following its entry in the cell. Thus we examined if the lifespan extension was dependent on this metabolic conversion. Using RNAi against the nematode’s only LDH orthologue *ldh-1*, we observed a loss of lifespan extension by lactate but not by pyruvate **(Supplementary Fig. 6g, h)**. This observation indicates that the oxidation of lactate to pyruvate is required to increase survival. Previous work identified *slcf-1* as an MCT orthologue in the nematode^55^. By knocking down the SLCF1 transporter using RNAi, we found that the lifespan extension by lactate or pyruvate was only partially affected **(Supplementary Fig. 6i)**, suggesting the existence of other MCTs or channels (e.g., pannexins) in nematodes, importing lactate into the cell.

We examined whether genes required for both stress resistance and longevity phenotypes could influence the expression of the ROS detoxifying reporter *gst-4::GFP*. Co-treatment of wild-type worms with NAC blocked the activation of *gst-4* induced by lactate or pyruvate supplementation **(Fig. 6i)**. Although mutation in *hif-1* surprisingly increased *gst-4* activity above control levels, both metabolites did not further increase the levels of GST-4::GFP **(Fig. 6j, Supplementary Fig. 7a, c)**. Likewise, a mutation in *pmk-3* **(Fig. 6k, Supplementary Fig. 7d-f)**, *ife-2* **(Fig. 6l, Supplementary Fig. 7g-i)** and *ire-1* **(Fig. 6m, Supplementary Fig. 7j-l)** reduced the basal activity of *gst-4*. Supplementation with lactate or pyruvate, although marginally increasing the reporter expression in *ife-2* and *ire-1* mutants, failed to restore the *gst-4* activity to wild-type levels. These observations strengthen the notion that lactate and pyruvate produce mild ROS elevations, which in turn trigger an increase in the expression of detoxifying mechanisms promoting stress resistance and longevity.

**Figure 7:**
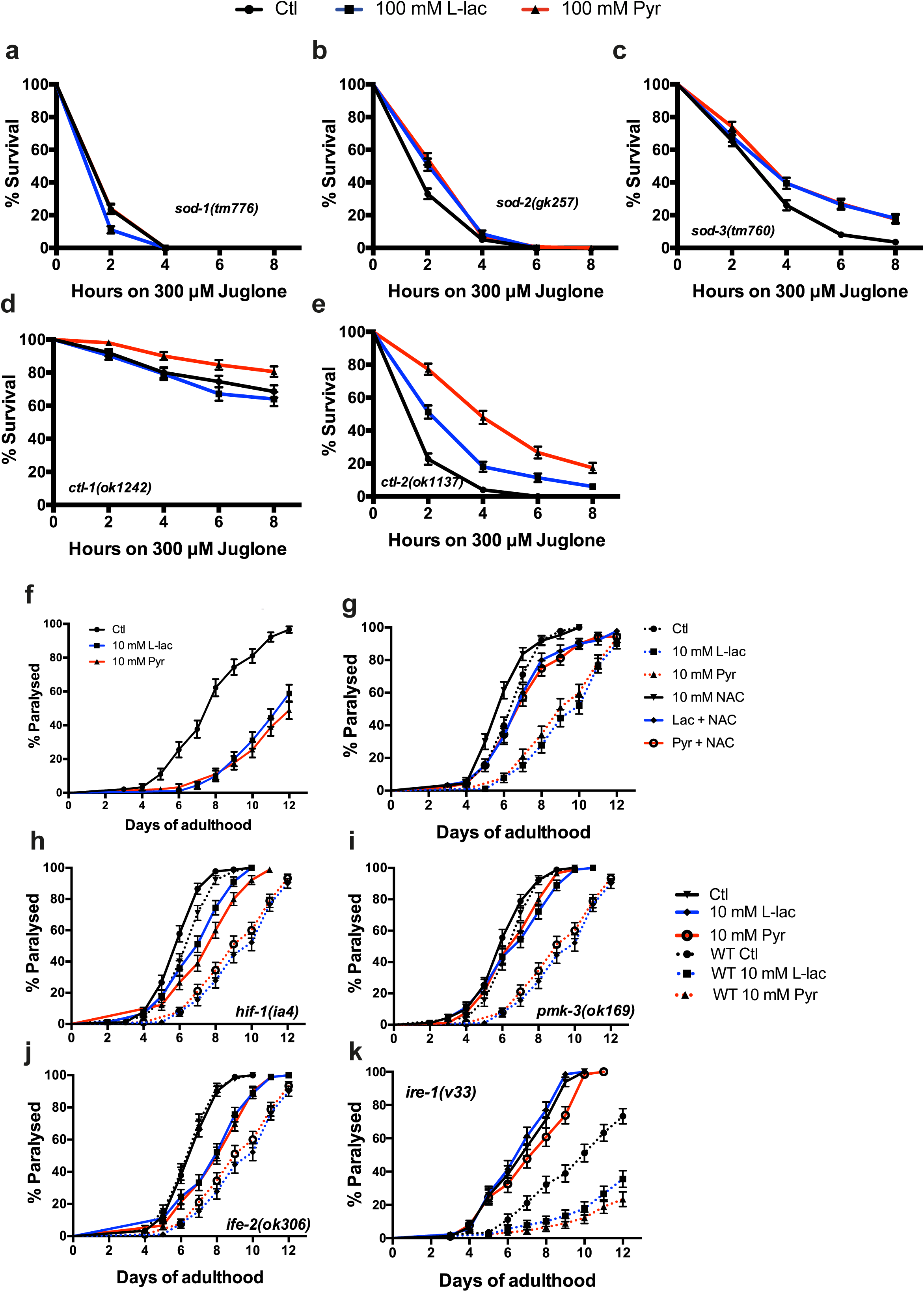
ROS detoxifying enzymes influence stress resistance and longevity induced by lactate and pyruvate. **a-c)** Survival curves for superoxide dismutase mutants treated with 300µM juglone and supplemented with 100 mM lactate or pyruvate. **d-e)** Survival curves for catalases mutants treated with 300µM juglone and supplemented with 100 mM lactate or pyruvate. **f-g)** Paralysis curves for *rgef-1p::Q67::CFP (Q67)* animals supplemented with 10 mM lactate or pyruvate **f)**. Q67 paralysis was assessed in the presence of 10 mM NAC **g) (h-k)** Paralysis curves for *rgef-1p::Q67::CFP (Q67)* animals supplemented with 10 mM lactate or pyruvate. Paralysis was assessed in mutant backgrounds for *hif-1* **h)**, *pmk-3* **i)**, *ife-2* **j)** and *ire-1* **k)** Statistical significance for the survival and paralysis curves was determined using the Log-Rank (Mantel-Cox) test. 90 animals were used per condition in each trial. Experiments were repeated at least three times. See supplementary table I for lifespan statistics.

### ROS detoxifying enzymes influence stress resistance by lactate and pyruvate

Given the findings that mild increases in ROS levels are involved in stress survival and longevity, we sought to investigate the dependence of ROS detoxifying enzymes on these phenotypes. We observed that *sod-1* and *sod-2* loss of function mutations prevented stress resistance by lactate or pyruvate, while *sod-3* mutants had no effects on lactate- or pyruvate-mediated protection **(Fig. 7a, c)**. Additionally, we also assessed the function of catalases. Loss of function mutants of *ctl-1* displayed a high stress resistance that was only increased by pyruvate, while *ctl-2* mutation did not influence lactate- or pyruvate-mediated stress resistance **(Fig. 7d, e)**.

### Lactate and pyruvate delay age-dependent neuronal dysfunction in a model of polyglutamine disease

As lactate and pyruvate promote stress resistance and longevity, we wondered whether both metabolites would delay neurodegeneration in a model of polyglutamine disease (*rgefp*::Q67::CFP, Q67). The pan-neuronal expression of polyglutamine tracts with 67 residues induces a late-onset paralysis. Upon supplementation with lactate and pyruvate, we observed a delayed age-dependent paralysis **(Fig. 7g)**. We examined the role of ROS signaling and found that this protection afforded by both lactate and pyruvate was lost when NAC was co-applied with the metabolites **(Fig. 7h)**. The protection against paralysis by lactate and pyruvate was also lost in *hif-1* and *pmk-3* mutants **(Fig. 7i, j),** while *ife-2* mutants partially blocked the protective effects by the metabolites **(Fig. 7k)**. Finally, mutation of *ire-1*, a UPR^ER^ element involved in stress resistance by lactate and pyruvate, accelerated the paralysis rate **(Fig. 7l) and** neither lactate nor pyruvate were able to reduce this increased toxicity.

## Discussion

In this study, we report a new mechanism underlying the role of lactate in cell survival. Lactate is known to reduce the toxicity of different cellular insults such as glutamate excitotoxicity or cerebral ischemia^8,9^ and promotes recovery through activation of PI3K/AKT and mTOR pathways. To provide insights into the molecular mechanisms of these positive effects of lactate, we analyzed how monocarboxylates influence cell survival against oxidative stress, in the context of aging and neurodegeneration. SH-SY5Y cells pre-treated with lactate showed increased survival under oxidative stress conditions. In line with previous findings^9^, transcriptome analysis of these cells revealed that lactate induces activation of PI3K/AKT and mTOR pathways. Interestingly, lactate also promotes the expression of DNAJ chaperones and unfolded protein response (UPR) genes **(Supplementary Fig. 2e-g)**, elements well-known to underlie the endoplasmic reticulum stress response^56^. Taken together, these observations indicate that lactate improves cell survival through an increase in ER protein homeostasis as well as via activation of the PI3K signaling pathway.

Lactate is at the center of a metabolic crossroad of glycolysis and oxidative metabolism. It was recently shown that lactate induces ROS^32,33^ a mandatory by-product of mitochondrial respiration and a signaling molecule^17^. We report here that lactate increases ROS in neuroblastoma and *C. elegans*, promoting cell survival through the mild activation of pro-oxidative mechanisms. These observations are in line with mitohormesis, a concept described in multiple organisms^18,57^ whereby a moderate increase in ROS production, in the low micromolar range **(Fig. 3f),** promotes stress resistance and longevity^20,58^. This hypothesis is supported by recent literature pointing to the signaling role of ROS^16,17^. Thus, results reported here **(Fig. 3 and 4)** suggest that lactate supports cell survival through a mild elevation in ROS levels that involves a boost in mitochondrial activity indicative of mitohormesis. Moreover, ROS increase PI3K, mTOR signaling as well as HIF1α activation *in vitro*^59^, suggesting that the mild pro-oxidative environment activates those pathways to promote cell survival.

NRF2/*skn-1* is a crucial transcription factor activated by oxidative stress^60^. The present results in cells and nematodes show increased activation of NRF2 and its downstream effector *gst-4*, respectively. ROS induction has to be tightly controlled to induce pro-survival outcomes^61^ and results reported here confirmed it. We observed that *pmk-3, ife-2* and *ire-1* loss of function all preventing normal activation of *gst-4*, did not support lactate- and pyruvate-mediated stress resistance **(Fig. 6J)**. However, *hif-1* mutants, although displaying an increased expression of *gst-4* compared to wild-type background, failed to respond to lactate- or pyruvate-increased stress resistance **(Fig. 6j).** Therefore, it appears that the activation window has to be controlled carefully to produce positive outcomes.

Our investigations in *C. elegans* also revealed that the concentration and the nature of the cellular stress have different outcomes, as shown by others^62,63^. Feeding nematodes with high concentrations (100 mM) of lactate or pyruvate increased resistance to oxidative stress and reduced lifespan while low concentrations (10 mM) increased lifespan but failed to induce stress resistance. This difference also translates into the genetic requirement underlying both protective phenomena. The IIS (*akt-1, age-1*), a branch of the UPR^ER^ (*ire-1),* and UPR^mt^ *(atfs-1)* **(Fig. 5)** promote stress resistance while lifespan extension relied on another branch of the UPR^ER^ *(atf-6)* and mitochondrial ETC (*nuo-6, isp-1*) to promote longevity. A low dose is likely to promote proteostasis^64^ to extend aging stages as observed in the longevity phenotype reported here **(Fig. 4a)**, whereas high doses produce a strong stimulation of stress relief mechanisms such as the UPR^mt^ that can increase stress resistance but are deleterious on the long term.

A question remains whether lactate supports cell survival by boosting metabolism through increased mitochondrial respiration and ATP production^5,9,65^ or through changes in the NADH/NAD+ redox ratio due to the conversion of lactate to pyruvate by the lactate dehydrogenase (LDH)^66^. Here, we found that lactate induced a strong resistance to oxidative stress and ROS induction *in vitro*, whereas pyruvate had a negligible impact in either of these processes. These observations are line with a protective mechanism relying on the production of NADH with the concomitant oxidation of lactate into pyruvate by LDH. However, findings in *C. elegans* differ significantly, as both lactate and pyruvate induce ROS, increasing stress resistance and promoting longevity. These data in the nematode suggest that the anti-aging effects of the monocarboxylates are mediated by ATP production. One explanation for this difference is that SH-SY5Y are cancer cells, which are known to rely mainly on glycolytic metabolism. Thus, the large amount of NADH produced by the metabolism of lactate by LDH may contribute to the stimulation of mitochondrial activity and mild elevation in ROS levels, a process not evoked by pyruvate.

### Conclusion and Perspective

In this report, we show that lactate promotes stress resistance *in vitro*, while both lactate and pyruvate promote stress resistance and longevity in *C. elegans.* It appears that in both models, this protective mechanism occurs through ROS signaling. It would be of interest to investigate whether this supporting mechanism is taking place between astrocytes and neurons in the mammalian brain^6^. Aerobic glycolysis^67^ and lactate release decrease during aging and in different neurodegenerative disorders^68^. Boosting the production and release of lactate by astrocytes or providing controlled chronic mild oxidative stress to neurons would be a promising therapeutic strategy to favor brain health.

## Methods

### Cell culture

SH-SY5Y neuroblastoma cells were grown in DMEM-F12 media (*Gibco - #11320-074*) supplemented with 10% FBS (*Gibco - #3000008085*) and 1% Pen/Strep mix (*Gibco - #15140122*), at 37°C with 5% CO2. Cells were maintained in T75 flasks, and media was changed every 2 days. For the experiments, cells were subcultured in different dish formats (6/24 wells plates) and used 36 hours after seeding. Sodium L-lactate (*Sigma - #71718*) and sodium pyruvate (*Sigma - #P2256*) were used in this study to investigate the role of both metabolites in cell survival.

For pharmacological treatments, inhibitors (**AR-C155858** – *Tocris #4960*, **LY294002** – *Sigma #L9908*, **Quercetin** – *Sigma #00200595*, **Cycloheximide** – *Sigma #C1988* and **N-acetylcysteine** – *Sigma #A7250*) were applied 15 min before lactate or pyruvate stimulation, and all treatments were conducted in DMEM-F12 media supplemented with 10% FBS.

### Cell viability assays

Cells were seeded 36h prior to assays to maximize their recovery and to allow a confluence of around 75-80% at the time of treatment. The day of the death induction, cells were treated with lactate or pyruvate for 6h in DMEM-F12 + 10% FBS. The media was then replaced with DMEM-F12 containing fresh H_2_O_2_ at a final concentration of 150µM. The next day the media was collected and cells trypsinized for analysis. The percentage of cells excluding the trypan blue stain was determined using a Countess II FL Automated Cell Counter (ThermoFisher) according to the user manual’s instructions.

For the MTT assay, MTT (3-[4,5-dimethylthiazol-2-yl]-2,5 diphenyl tetrazolium bromide) was added to cell media at a final concentration of 0.2 mg/ml. Cells were treated for 2h to allow the transformation to formazan. After incubation, media was replaced by DMSO to dissolve the cristals and absorbance was measure at λ=570 nm.

### ROS detection

Cells were incubated in under different conditions: 1) 6h with 20mM lactate in the presence or absence of 0.1 mM N-acetylcysteine or 2) with 150µM H_2_O_2_ for 30 min or 3) 6h with 20 mM pyruvate. After the initial treatments, 100 µM 2’,7’-dichlorodihydrofluorescein diacetate (H_2_DCFDA) was added to the media for 30 min. Cells were washed in HBSS media (136 mM NaCl, 3 mM KCl, 1,25 mM CaCl_2_, 1,25 mM MgSO_4_, 10 mM HEPES and 2 mM D-glucose) and ROS levels were measured.

Nematodes were treated with 100µM H_2_DCFDA for a 1h, washed 3 times with M9 buffer (3g/L KH_2_PO_4_, 6g/L Na_2_HPO_4_, 5g/L NaCl and 1mM MgSO_4_) and scored on a 2% agarose pad with an M9 buffer with 5 mM Levamisole. Both cells and worms ROS levels were measured using a Zeiss LSM780 and AxioExaminer at λ=588 nm.

### JC1 staining

Cells were incubated in under different conditions: 6h with 20mM lactate or with 150µM H_2_O_2_ for 30 min. After the initial treatments, 2.5 µM of JC-1 staining was added to the media for 10 min. Cells were washed in HBSS media (136 mM NaCl, 3 mM KCl, 1,25 mM CaCl_2_, 1,25 mM MgSO_4_, 10 mM HEPES and 2 mM D-glucose) and fluorescence levels were measured using a Zeiss LSM780 and AxioExaminer at λ=588 nm and 561 nm.

### RNA extraction

Total RNA from SH-SY5Y cells was isolated RNeasy plus mini kits (Qiagen) following the manufacturer’s instruction.

### RNA sequencing methods

Concentration, purity and integrity of the RNA extracted from the neuroblastoma cells were assessed with a NanoDrop spectrophotometer (NanoDrop 2000, ThermoFisher Scientific), and a 2100 Bioanalyzer (Agilent).

Total RNA with an RNA Integrity Number above 9.5 was used to construct libraries using the TruSeq Stranded mRNA Sample Kit (Illumina) following the protocol’s instructions. Briefly, mRNA was enriched using oligo dT-attached magnetic beads, fragmented, and converted into cDNA. Fragments of cDNA went through an end repair process, 3’ ends were adenylated, universal bar-coded adapters were ligated, and cDNA fragments were amplified by PCR to yield the final libraries. The sequencing libraries were evaluated using a 2100 Bioanalyzer (Agilent). Paired-end read (2 x 150 bp) multiplex sequencing from pooled libraries was performed on an Illumina HiSeq 4000 machine at the KAUST Bioscience Core Labs. An average of 40-50 million reads was obtained for each sample. Sequencing data have been deposited in the NCBI SRA database under the project accession number PRJNA510906.

Raw read quality was evaluated with the FastQC tool (https://www.bioinformatics.babraham.ac.uk/projects/fastqc/). Low-quality reads were filtered out and adapter sequences trimmed using Trimmomatic version 0.36 ^69^ with the following parameters: ILLUMINACLIP/TruSeq3-PE-2.fa:2:30:10, LEADING:3, TRAILING:3, SLIDINGWINDOW:4:15, MINLEN:36. Reads from each sample replicate were mapped to the mouse reference genome (Ensembl, release 91) using STAR version 2.6.0a ^70^ with default parameters except for outFilterMultimapNmax set to 1 (using Hisat2 version 2.1.0 ^71^ with default parameters except for k set to 1). Mapped reads for protein-expressing genes were summarized with the featureCounts program (Subread package, version 1.5.2, ^72^), and the differential expression analysis was performed according to using the Bioconductor package DESeq2 ^73^ in the R programming environment. To minimise background noise and to focus on more significant genes in term of biological impact, we removed genes with very low expression levels, excluding genes that failed to total an average count above 10 in any conditions. Differentially expressed genes (DEG) were considered in pairwise comparisons with a threshold including a fold change expression ≥ 1.5 and q-value (or False Discovery Rate, FDR) < 0.05. To obtain a functional representation of the lists of DEG, we performed gene ontology (GO) and Biocarta and KEGG pathways enrichment analyses using the online database and tool DAVID (version 6.8, https://david.ncifcrf.gov).

### C. elegans

Standard methods of culturing and handling worms were used. Worms were maintained on standard NGM plates streaked with OP50 Escherichia coli, and all strains were scored at 20 °C unless indicated. Mutant strains were obtained from the *C. elegans* Genetics Center (University of Minnesota, Minneapolis, MN, USA). Mutants or transgenic worms were verified by visible phenotypes, PCR analysis for deletion mutants, sequencing for point mutations, or a combination thereof. Deletion mutants were out-crossed a minimum of three times to wild-type worms before use. The strains list can be found in **Supp. Table II**.

### The measure of ageing phenotypes in C. elegans

#### Juglone (oxidative stress) test

Worms were grown on NGM (OP50-1) in the presence or absence of 100 mM lactate or pyruvate until and then transferred to NGM containing 300µM juglone (5-Hydroxy-1,4-naphthoquinone, Sigma). Juglone was dissolved in 96% ethanol.

#### Paralysis test

Worms were grown on NGM (OP50-1) in the presence or absence 10mM lactate or pyruvate. Experiments were conducted at 20°C, in 3 independent trials using 90 animals for each trial. Animals were respectively scored as paralysed or dead (excluded from the analysis when locomotion or head movement was not observed following nose prodding using a platinum wire.

#### Lifespan assay

Nematodes were grown on NGM supplemented or not with lactate or pyruvate. Adult day 1 one animals, 30 per plates in triplicates, were transferred onto NGM supplemented with 20µM FUdR, and lifespan was assessed at 20°C. Animals were declared dead if they failed to respond to tactile or heat stimuli. For dead bacteria experiments, OP50-1 bacteria were heat-killed by incubating the culture at 80°C for 4h. For the RNAi experiments, animals were fed with E.coli (HT115) containing an empty vector, or either *ldh-1*(F13D12.2), *skn-1*(TE19E7.2) and *slcf-1*(F59F5.1).

#### Immunoblotting

SH-SY5Y cells were harvested from 6 wells plate and washed with cold 1X PBS. *C. elegans* were collected in M9 buffer, and the pellets were quickly frozen at −80°C overnight. Both tissues were lysed using RIPA buffer (150 mM NaCl, 50 mM Tris pH 7.4, 1% Triton X-100, 0.1% SDS, 1% sodium deoxycholate) containing 1X Protease and Phosphatase Inhibitor Cocktail (*ThermoFisher #78440*). Protein concentration was measured using Bicinchoninic assay (BCA – Thermofisher). Protein extracts were then loaded on 10% SDS-page acrylamide gels and transferred on PVDF membrane *(Millipore #IPVH00010)* overnight. The membranes were blocked using in PBS containing 0.1%Tween and 5% milk for at least 30 min. Primary antibodies were applied overnight at 4°C, and subsequently, HRP-conjugated secondary antibodies were incubated for 1h before visualisation using ECL (*ThermoFisher #34096*)

### Quantification and statistical analysis

GraphPad Prism 7 was used for statistical analysis and ImageJ to quantify the images and the. The Log-rank (Mantel-Cox) method was used to compare survival curves. Statistical analyses for all of the data except for lifespan was carried out using or ANOVA or Student’s *t*-test (unpaired, two-tailed) Experiments were performed in triplicate. A p-value <0.05 was considered statistically significant. Data are expressed as mean ± SEM unless otherwise indicated.

## Acknowledgments

We thank Magdalena Julkowska, Nadia Steiner and Lorène Mottier for their technical assistance. We want to thank Dr. J. Alex Parker initial support in the project and RNAi clones and Dr. Christian Frøkjær-Jensen for critical reading of the manuscript. Thanks to the *C. elegans* Genetic Center for the different strains used in this study.

Support for this study was provided by King Abdullah University of Science and Technology.

## Author contributions

AT performed the experiments. SAM helped with data acquisition. AT, HF and PJM designed the experiments, commented on the results and reviewed the manuscript.

## Materials & Correspondence

Requests and correspondence: Arnaud Tauffenberger or Pierre Magistretti

