## Supplementary Information for "Lactate and pyruvate promote cellular stress resistance and longevity through ROS signaling"

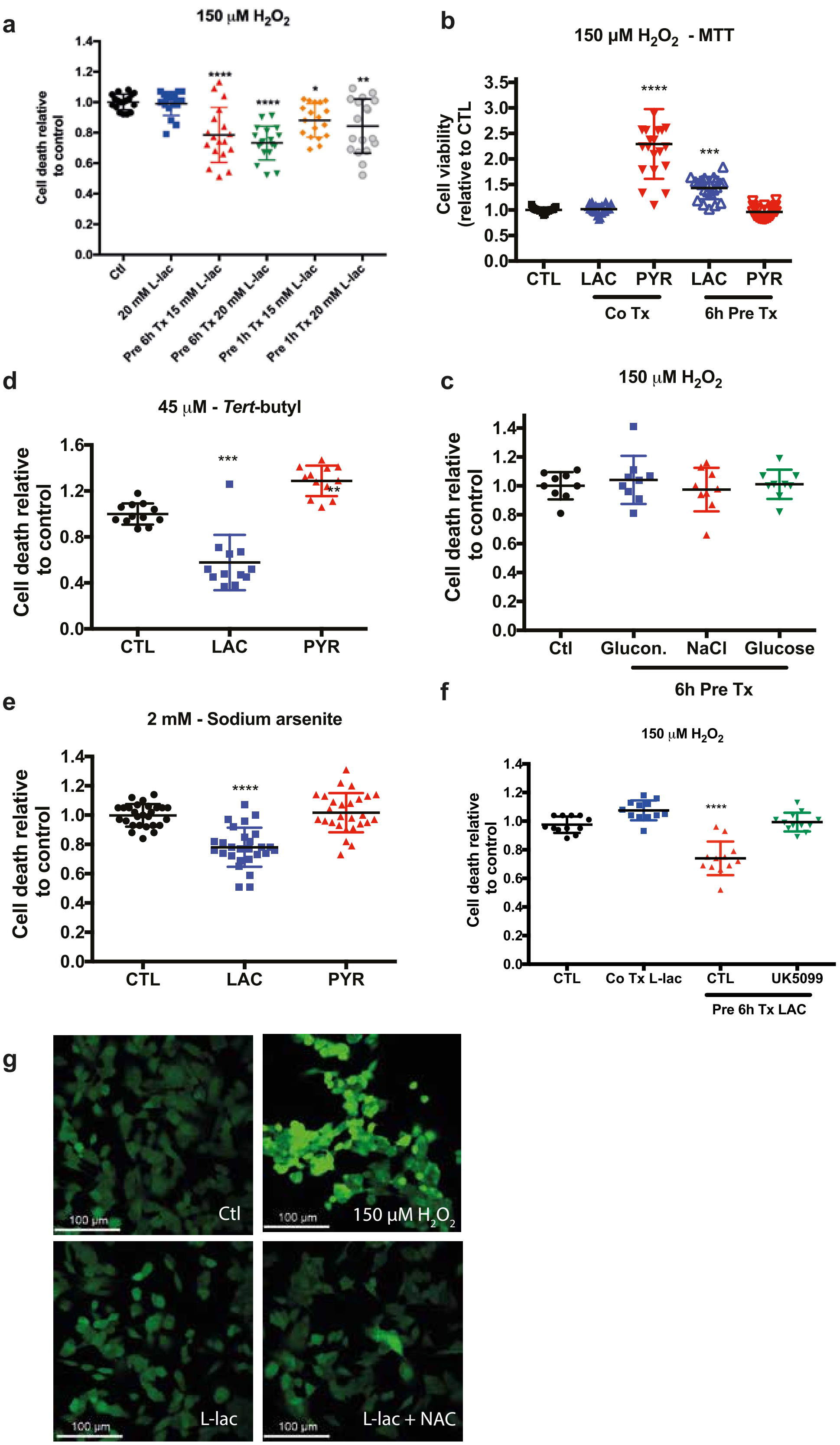
a
150 μM H2O2
Cell death relative to control
Ctl
20 mM L-lac
Pre 6h Tx 15 mM L-lac
Pre 6h Tx 20 mM L-lac
Pre 1h Tx 15 mM L-lac
Pre 1h Tx 20 mM L-lac
b
150 μM H2O2 - MTT
Cell viability (relative to CTL
CTL
LAC
PYR
LAC
PYR
Co Tx
6h Pre Tx
d
45 μM - Tert-butyl
Cell death relative to control
CTL
LAC
PYR
c
150 μM H2O2
Cell death relative to control
Ctl
Glucon.
NaCl
Glucose
6h Pre Tx
e
2 mM - Sodium arsenite
Cell death relative to control
CTL
LAC
PYR
f
150 μM H2O2
Cell death relative to control
CTL
Co Tx L-lac
CTL
UK5099
Pre 6h Tx LAC
g
100 μm
Ctl
150 μM H2O2
L-lac
L-lac + NAC

**Supplementary Figure 1: Characterization of lactate protective response.** ***(Related to Figure 1)***

**a)** A measure of cell death after 150 μM $H_2O_2$ treatment upon pre-treatment with lactate (15mM -20mM) for different durations (1h – 6h).

**b)** MTT measure of cell death upon pre-treatment with lactate (20 mM – 6h).

**c)** A measure of cell death after 150 μM $H_2O_2$ treatment upon pre-treatment with D-glucose, NaCl or Na-gluconate (all 20mM).

**d-e)** Measure of cell death upon pre-treatment with lactate (20 mM – 6h) after treatment with 45 μM tert-butyl **d)** or 2mM Sodium Arsenite **e)**.

**f)** The measure of cell death after 150 μM $H_2O_2$ and upon pre-treatment with lactate (20 mM – 6h) using MCT blocker UK5099.

**g)** Confocal images of SH-SY5Y cells stained with $H_2$DCFDA (50 μM). Cells were treated with 150 μM $H_2O_2$ for 30 min or with 20mM lactate for 6h.

Each data point represents one measurement. Bars are the average ± SEM. All calculations for statistical significance were completed using a non-parametric, one-way ANOVA with multiple comparisons, n= 12-15 from 4-5 independent experiments. *$p<0.05$, **$p<0.01$, ***$p<0.001$, ****$p<0.0001$

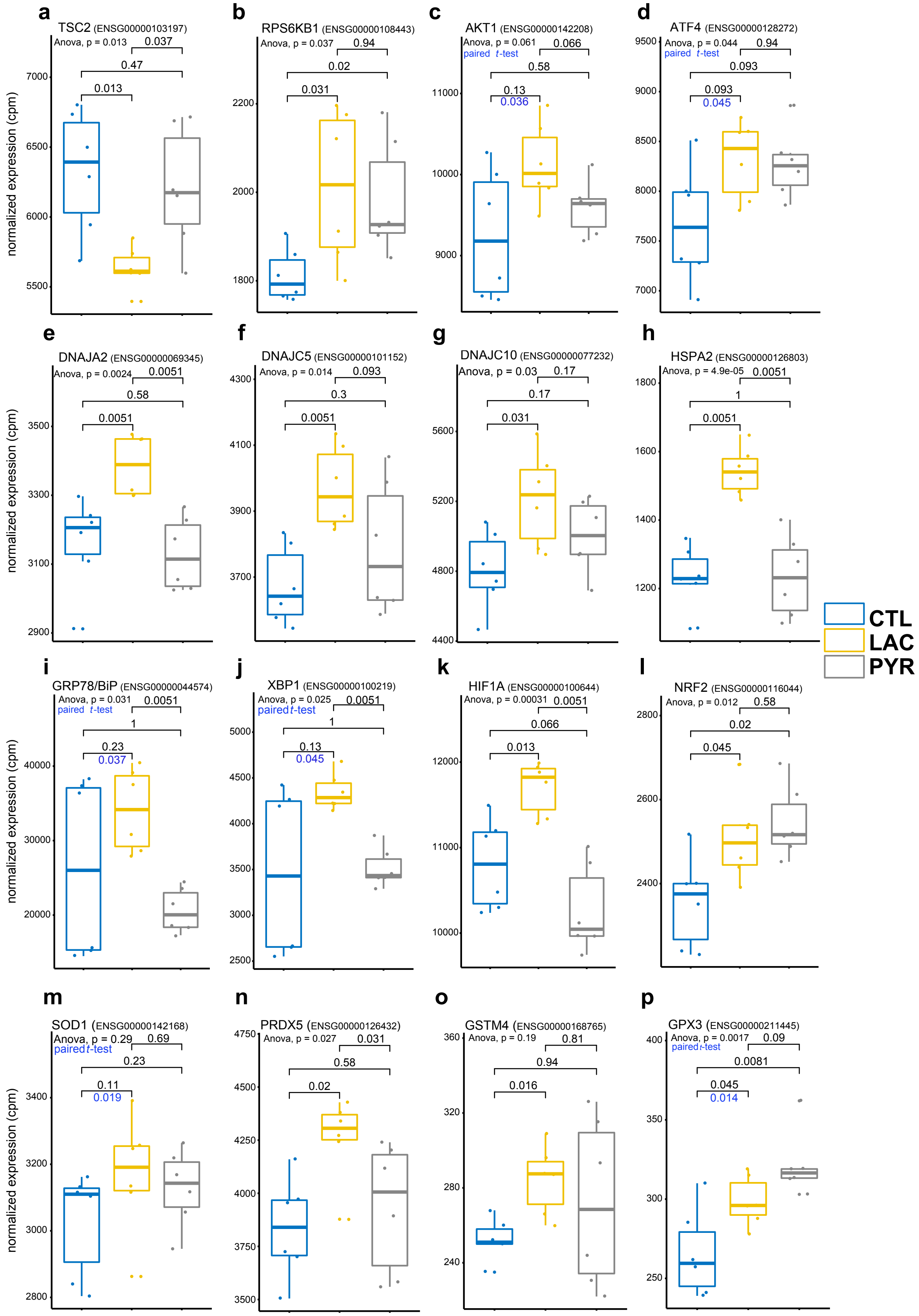

a
TSC2 (ENSG00000103197)
Anova, p = 0.013
0.037
0.47
0.013
b
RPS6KB1 (ENSG00000108443)
Anova, p = 0.037
0.94
0.02
0.031
c
AKT1 (ENSG00000142208)
Anova, p = 0.061
paired t-test
0.066
0.58
0.13
0.036
d
ATF4 (ENSG00000128272)
Anova, p = 0.044
paired t-test
0.94
0.093
0.093
0.045
e
DNAJA2 (ENSG00000069345)
Anova, p = 0.0024
0.0051
0.58
0.0051
f
DNAJC5 (ENSG00000101152)
Anova, p = 0.014
0.093
0.3
0.0051
g
DNAJC10 (ENSG00000077232)
Anova, p = 0.03
0.17
0.17
0.031
h
HSPA2 (ENSG00000126803)
Anova, p = 4.9e-05
0.0051
1
0.0051
i
GRP78/BiP (ENSG00000044574)
Anova, p = 0.031
paired t-test
0.0051
1
0.23
0.037
j
XBP1 (ENSG00000100219)
Anova, p = 0.025
paired t-test
0.0051
1
0.13
0.045
k
HIF1A (ENSG00000100644)
Anova, p = 0.00031
0.0051
0.066
0.013
l
NRF2 (ENSG00000116044)
Anova, p = 0.012
0.58
0.02
0.045
m
SOD1 (ENSG00000142168)
Anova, p = 0.29
paired t-test
0.69
0.23
0.11
0.019
n
PRDX5 (ENSG00000126432)
Anova, p = 0.027
0.031
0.58
0.02
o
GSTM4 (ENSG00000168765)
Anova, p = 0.19
0.81
0.94
0.016
p
GPX3 (ENSG00000211445)
Anova, p = 0.0017
paired t-test
0.09
0.0081
0.045
0.014
normalized expression (cpm)
CTL
LAC
PYR

**Supplementary Figure 2: Expression levels of candidates identified in the RNAseq analysis of SH-SY5Y cells after lactate treatment**. ***(Related to Figure 2)***

**a-d)** Transcript levels of mTOR and PI3K related candidates; **a)** TSC2, **b)** RPS6KB1, **c)** AKT1 and **d)** ATF4.

**e-k)** Transcript levels for ER processing related candidates; **e)** DNAJA2, **f)** DNAJC5, **g)** DNAJC10, **h)** HSPA2 **i)** GRP78/BiP, and **j)** XBP1.

**k-p)** Transcripts levels of candidates related to ROS induction and detoxification; **k)** HIF1a, **l)** NRF2, **m)** SOD1, **n)** PRDX53, **o)** GSTM4 and **p)** GPX3.

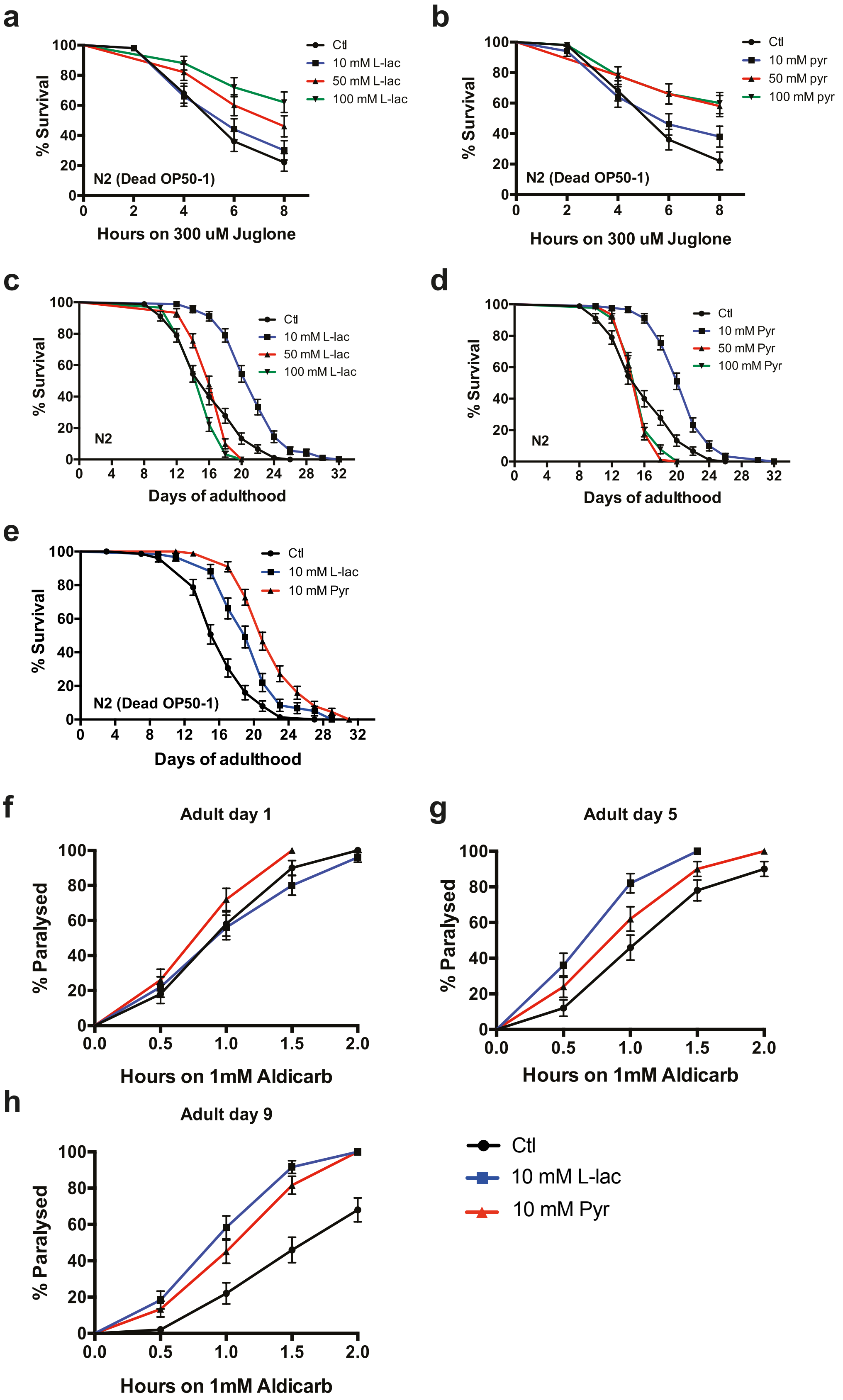

a
% Survival
Hours on 300 uM Juglone
N2 (Dead OP50-1)
Ctl
10 mM L-lac
50 mM L-lac
100 mM L-lac
b
% Survival
Hours on 300 uM Juglone
N2 (Dead OP50-1)
Ctl
10 mM pyr
50 mM pyr
100 mM pyr
c
% Survival
Days of adulthood
N2
Ctl
10 mM L-lac
50 mM L-lac
100 mM L-lac
d
% Survival
Days of adulthood
N2
Ctl
10 mM Pyr
50 mM Pyr
100 mM Pyr
e
% Survival
Days of adulthood
N2 (Dead OP50-1)
Ctl
10 mM L-lac
10 mM Pyr
f
Adult day 1
% Paralysed
Hours on 1mM Aldicarb
g
Adult day 5
% Paralysed
Hours on 1mM Aldicarb
h
Adult day 9
% Paralysed
Hours on 1mM Aldicarb
Ctl
10 mM L-lac
10 mM Pyr

**Supplementary Figure 3: Characterization of longevity phenotypes induced by lactate and pyruvate.** ***(Related to figure 4)***

**a-b)** Survival curves of wild-type (N2) animals treated with 300μM juglone and supplemented with 10, 50 and 100 mM lactate or pyruvate before the test. Worms were grown on dead OP50-1.

**c-d)** Lifespan curves of wild-type (N2) animals treated with 10, 50 and 100 mM lactate or pyruvate. **($p<0.0001$)**.

**e)** Lifespan curves of wild-type (N2) animals treated with 10 mM lactate or pyruvate. Worms were grown and tested on dead OP50-1. **($p<0.0001$)**

**g-i)** Paralysis curves of wild-type (N2) animals treated with 1 mM aldicarb after dietary supplementation with 10 mM lactate or pyruvate. Animals were tested adult day 1 **g)**, day 5 **h)** and day 9 **i)**. ($p<0.01$ **Ctl vs Pyr Ad5**; $p<0.0001$ **Ctl vs L-lac Ad5 and Ad9**; $p<0.0001$ **Ctl vs Pyr Ad9**)

Survival curves were generated and compared using the Log-rank (Mantel-Cox) test, and 90-100 animals were tested per genotype and repeated at least three times. See supplementary table I for lifespan statistics.

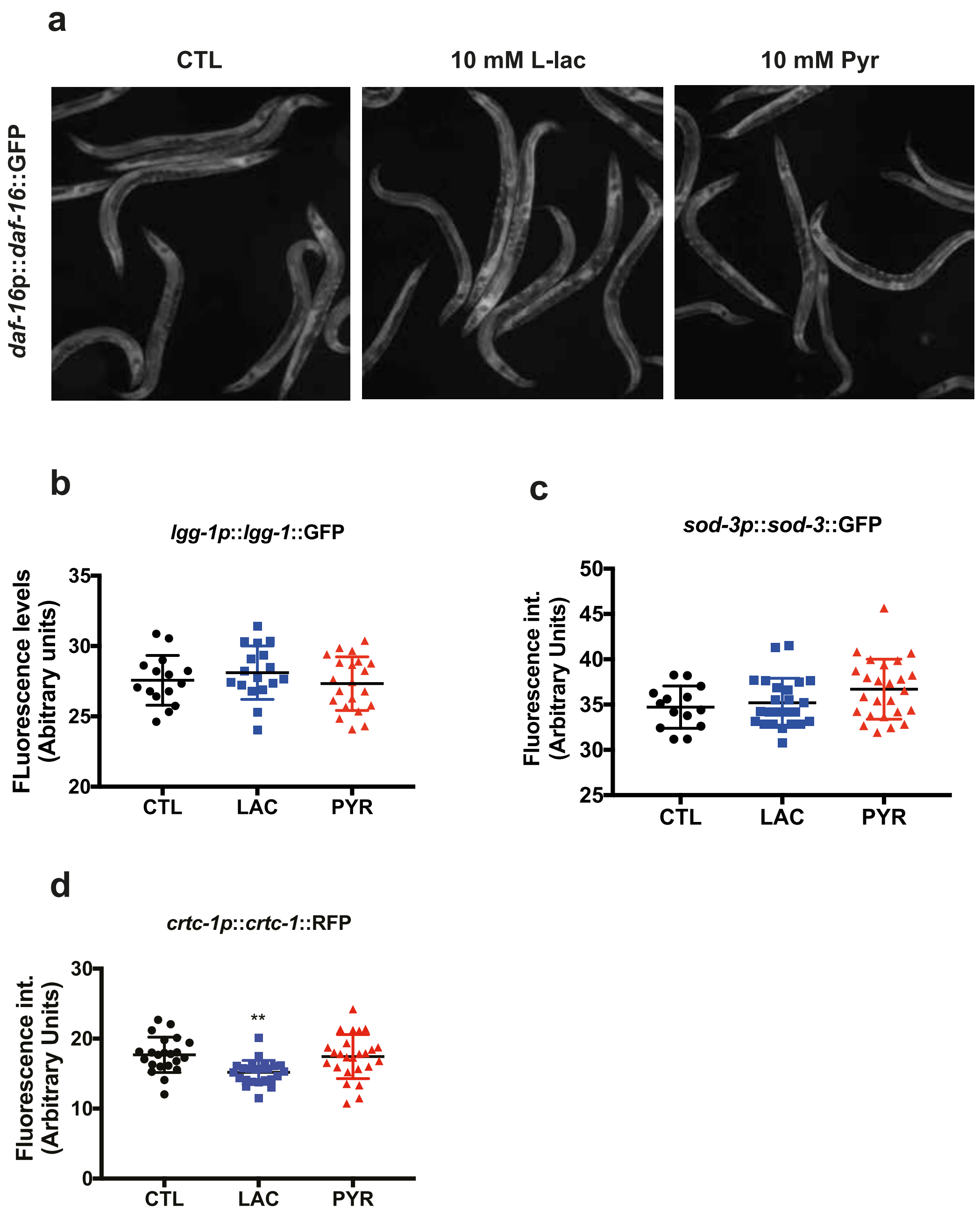


**Supplementary Figure 4: lactate and pyruvate do not increase the expression of stress-related response reporter.** ***(Related to Figure 4)***

**a-d)** Measure of expression for translational reporter of *daf-16* **a)**, *lgg-1* **b)**, *sod-3* **c)** and *crtc-1* **d)** after dietary supplementation with 10 mM lactate or pyruvate. **(p<0.01)

Each data point represents one measurement +/- SEM. All calculations for statistical significance were completed using a non-parametric, one-way anova with multiple comparisons, n= 12-15 from 4-6 independent experiments.

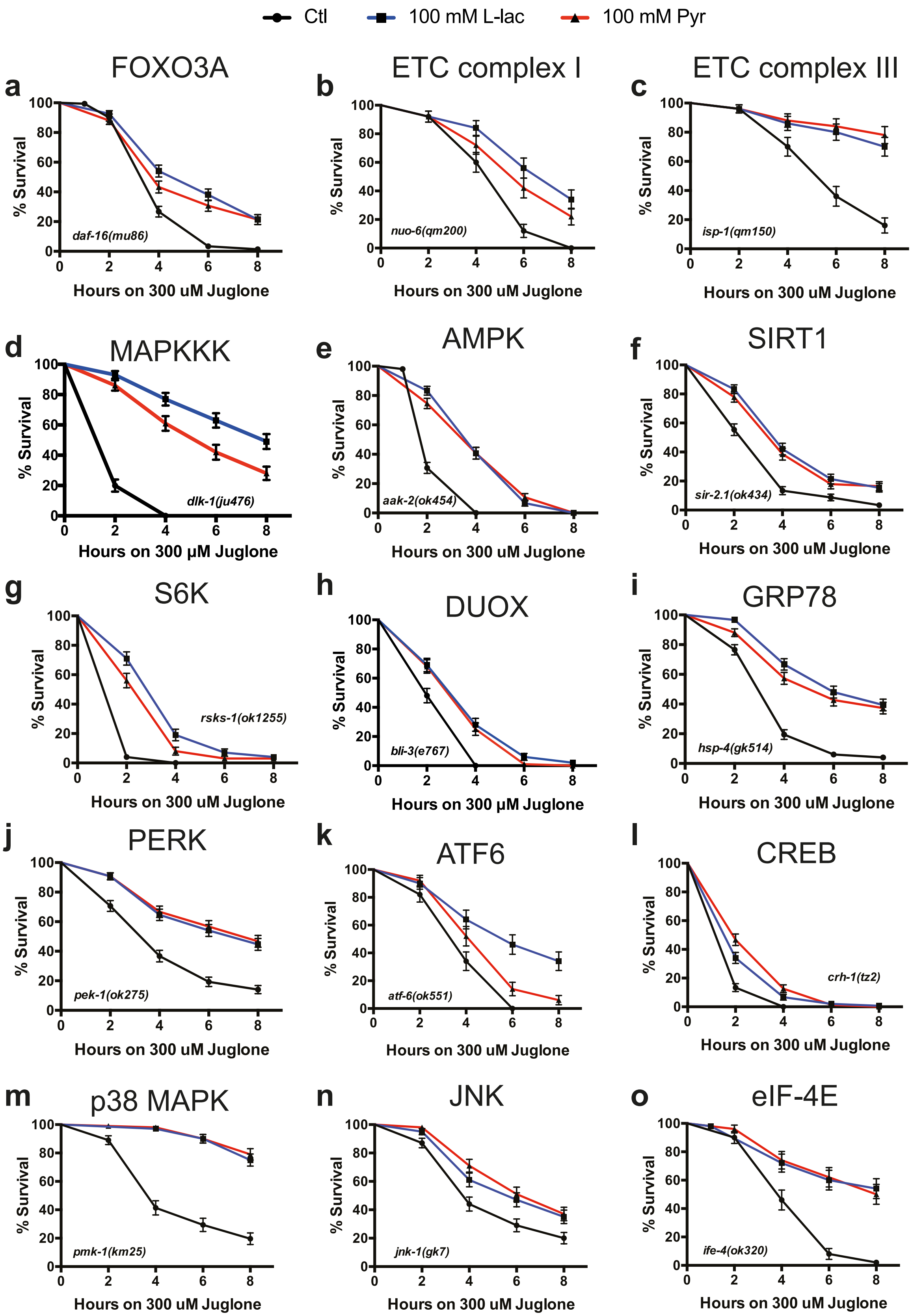

Ctl
100 mM L-lac
100 mM Pyr
a
FOXO3A
daf-16(mu86)
b
ETC complex I
nuo-6(qm200)
c
ETC complex III
isp-1(qm150)
d
MAPKKK
dlk-1(ju476)
e
AMPK
aak-2(ok454)
f
SIRT1
sir-2.1(ok434)
g
S6K
rsks-1(ok1255)
h
DUOX
bli-3(e767)
i
GRP78
hsp-4(gk514)
j
PERK
pek-1(ok275)
k
ATF6
atf-6(ok551)
l
CREB
crh-1(tz2)
m
p38 MAPK
pmk-1(km25)
n
JNK
jnk-1(gk7)
o
eIF-4E
ife-4(ok320)
% Survival
Hours on 300 uM Juglone
Hours on 300 μM Juglone

**Supplementary Figure 5: Different metabolic sensitive pathways are not required for lactate and pyruvate stress resistance.** ***(Related to figure 5)***

**a-o)** Survival curves of mutant nematodes treated 300µM juglone and supplemented with 100mM lactate or pyruvate. Mutated genes are indicated as an inset in the lower left or right corner of each graph, and mammalian orthologs are indicated above each graph.

Survival curves were generated and compared using the Log-rank (Mantel-Cox) test, and 90 animals were tested per genotype and repeated at least three times.

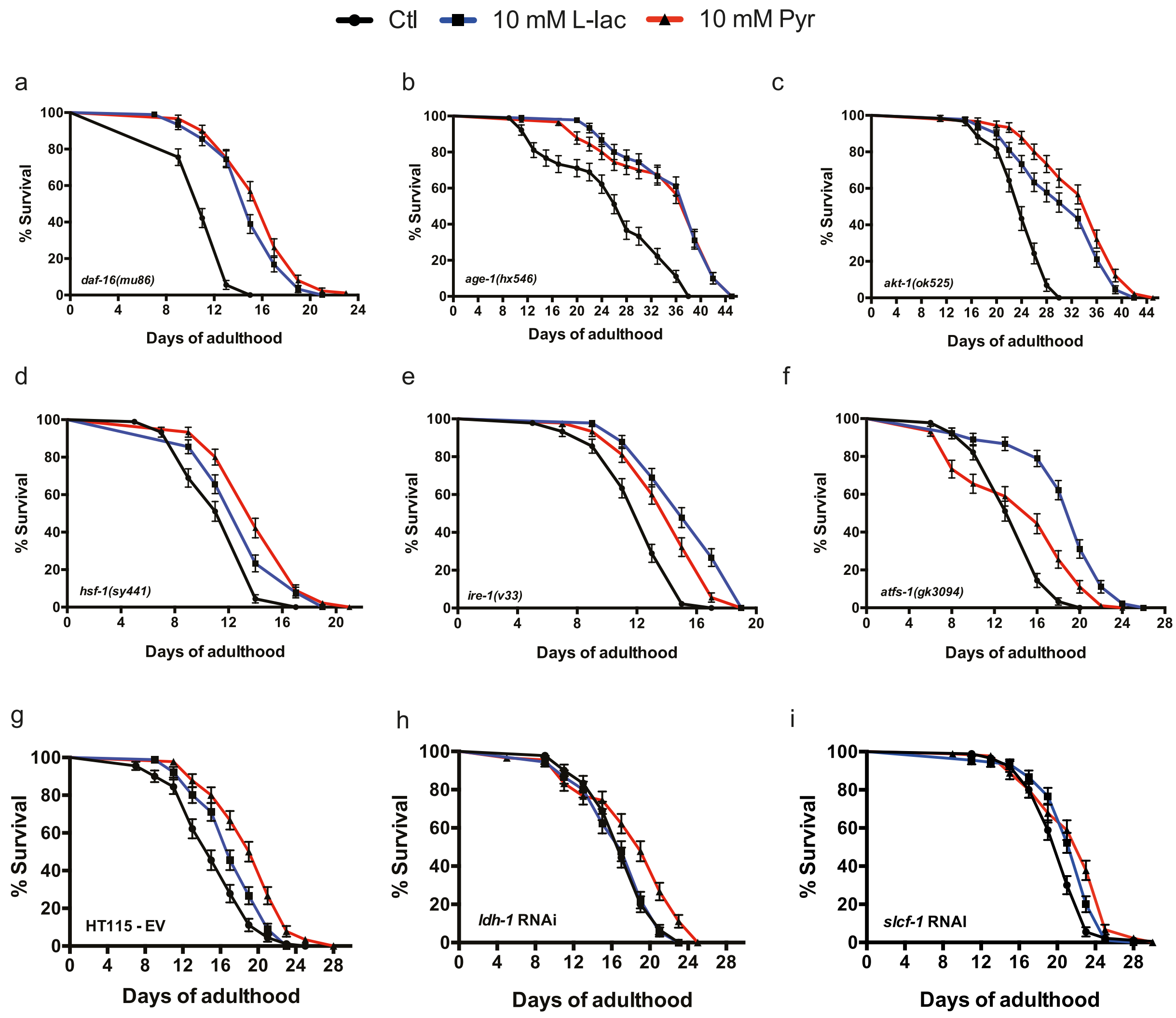


**Supplementary Figure 6: Characterization of the cellular pathways not involved in lactate and pyruvate longevity phenotype.** ***(Related to Figure 6)***

**a-c)** Lifespan curves of mutants animals supplemented with 10 mM lactate or pyruvate. IIS mutants for: **a)**, FOXO3A/*daf-16* **b)**, PI3K/*age-1* **c)** and AKT1/*akt-1*.

**d)** Lifespan curves of HSF1/*hsf-1* mutant supplemented with 10 mM lactate or pyruvate.

**e)** Lifespan curve of $UPR^{ER}$ kinase IRE1/*ire-1* mutant supplemented with 10 mM lactate or pyruvate.

**f)** Lifespan curve of $UPR^{mt}$ transcription factor *atfs-1* mutant supplemented with 10 mM lactate or pyruvate.

**g-i)** Lifespan curve of wild-type animals (N2) treatment with RNAi; Empty vector (EV) **g)**, *ldh-1* RNAi **h)** and *slcf-1* RNAi **i)** and supplemented with 10 mM lactate or pyruvate.

Survival curves were generated and compared using the Log-rank (Mantel-Cox) test, and 90 animals were tested per genotype and repeated at least three times. See supplementary table I for lifespan statistics.

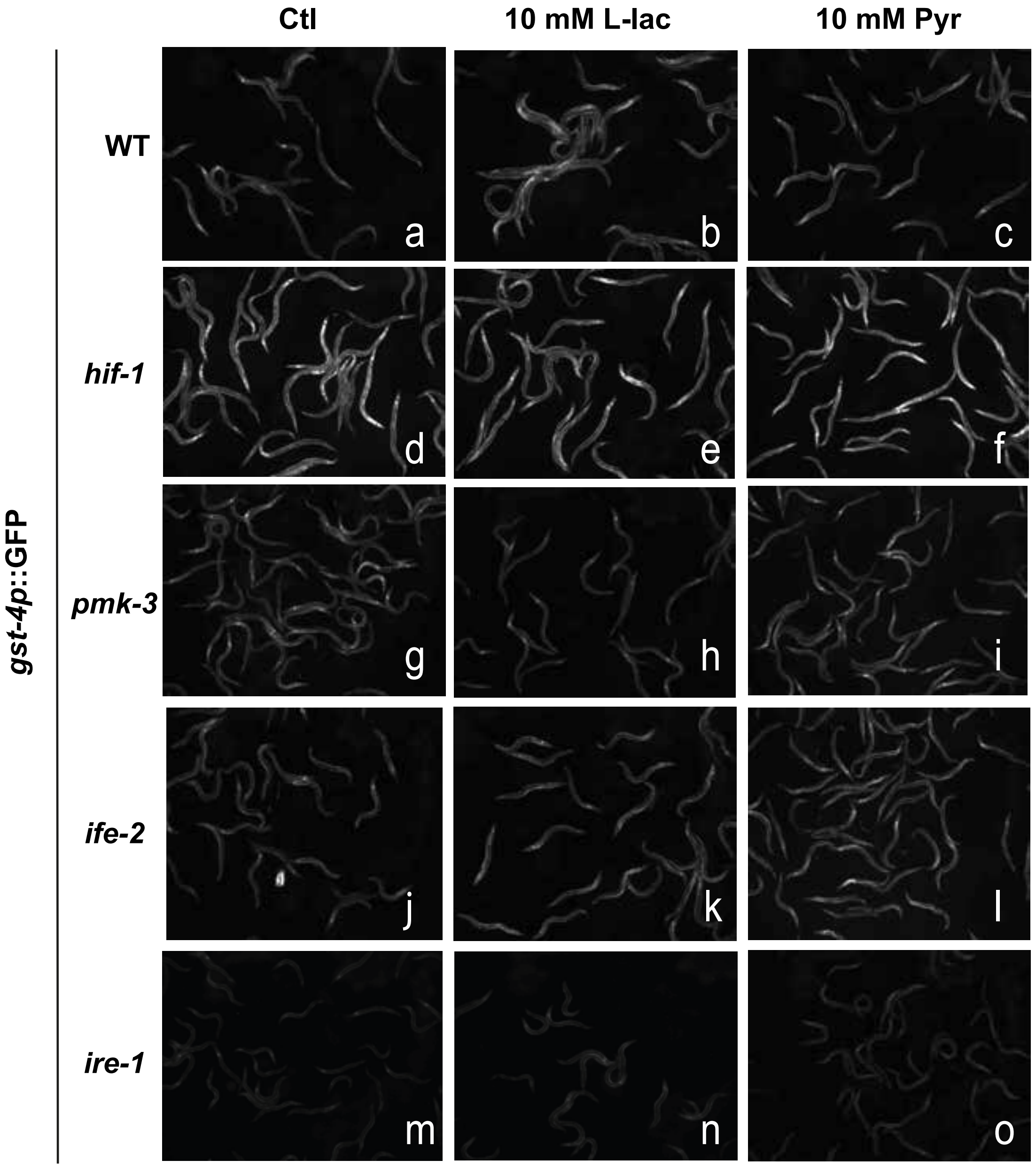


**Supplementary figure 7: Imaging of *gst-4p*::GFP animals with different mutant backgrounds, upon lactate and pyruvate treatment**. ***(Related to Figure 6)***

**a-c)** Wild-type animals imaging upon 10 mM lactate or pyruvate diet supplementation

**d-f)** Animals with *hif-1* mutant background imaging upon 10 mM lactate or pyruvate diet supplementation.

**g-i)** Animals with *pmk-3* mutant background imaging upon 10 mM lactate or pyruvate diet supplementation

**j-l)** Animals with *ife-2* mutant background imaging upon 10 mM lactate or pyruvate diet supplementation

**m-o)** Animals with *ire-1* mutant background imaging upon 10 mM lactate or pyruvate diet supplementation.

| Figure | Genotype | Mean lifespan | Maximum lifespan | P value | Numbers of animals |
|---|---|---|---|---|---|
| **4** | N2 | 16 | 26 | | 270 |
| | +10 mM L-lactate | 22 | 32 | <0.0001 | 270 |
| | +10 mM Pyruvate | 22 | 32 | <0.0001 | 270 |
| **6** | N2 | 14 | 26 | | 180 |
| | +10 mM L-lactate | 18 | 28 | <0.0001 | 180 |
| | +10 mM pyruvate | 18 | 30 | <0.0001 | 180 |
| | *hif-1(ia4)* | 10 | 18 | | 180 |
| | +10 mM L-lactate | 12 | 18 | n.s. 0.0636 | 180 |
| | +10 mM pyruvate | 12 | 18 | n.s. 0.0922 | 180 |
| | *pmk-3(ok169)* | 12 | 18 | | 180 |
| | +10 mM L-lactate | 13,5 | 20 | n.s. 0.4600 | 180 |
| | +10 mM pyruvate | 15 | 18 | n.s. 0.6157 | 180 |
| | *ife-2(ok3206)* | 16 | 25 | | 180 |
| | +10 mM L-lactate | 16 | 23 | n.s. 0.9013 | 180 |
| | +10 mM pyruvate | 16 | 25 | n.s. 0.6244 | 180 |
| | daf-2(e1370) | 39 | 53 | | 180 |
| | +10 mM L-lactate | 29 | 45 | <0.0001 | 180 |
| | +10 mM pyruvate | 31 | 43 | <0.0001 | 180 |
| | 10 mM NAC | 15 | 22 | | 180 |
| | +10 mM L-lactate | 17 | 22 | n.s. 0.0645 | 180 |
| | +10 mM pyruvate | 16 | 22 | n.s. 0.1772 | 180 |
| | *atf-6(ok551)* | 22 | 30 | | 180 |
| | +10 mM L-lactate | 19 | 24 | <0.0001 | 180 |
| | +10 mM pyruvate | 19 | 26 | 0.0003 | 180 |
| | *isp-1(qm150)* | 26 | 42 | | 180 |
| | +10 mM L-lactate | 25 | 40 | 0,0105 | 180 |
| | +10 mM pyruvate | 27 | 42 | n.s. 0.6080 | 180 |
| **Supp. 3** | **N2** | 16 | 26 | | 270 |
| | + 10 mM L-lactate | 22 | 32 | <0.0001 | 270 |
| | + 50 mM L-lactate | 16 | 20 | n.s. 0.4197 | 270 |
| | + 100 mM L-lactate | 16 | 20 | n.s. 0.0019 | 270 |
| | **N2** | 16 | 26 | | 270 |
| | + 10 mM pyruvate | 22 | 32 | <0.0001 | 270 |
| | + 50 mM pyruvate | 16 | 20 | 0.0068 | 270 |
| | + 100 mM pyruvate | 16 | 20 | 0.0275 | 270 |
| | **N2** | 17 | 27 | | 90 |
| | +10 mM L-lactate (Dead OP50-1) | 19 | 29 | <0.0001 | 90 |
| | +10 mM pyruvate (Dead OP50-1) | 21 | 31 | <0.0001 | 90 |
| **Supp. 6** | ***daf-16(mu86)*** | 11 | 15 | | 90 |
| | +10 mM L-lactate | 15 | 21 | <0.0001 | 90 |
| | +10 mM pyruvate | 17 | 23 | <0.0001 | 90 |

| | | | | |
|---|---|---|---|---|
| ***age-1(hx546)*** | 28 | 38 | | 90 |
| +10 mM L-lactate | 39 | 45 | <0.0001 | 90 |
| +10 mM pyruvate | 39 | 45 | <0.0001 | 90 |
| ***akt-1(ok525)*** | 24 | 30 | | 90 |
| +10 mM L-lactate | 33 | 42 | <0.0001 | 90 |
| +10 mM pyruvate | 36 | 45 | <0.0001 | 90 |
| *hsf-1(sy441)* | 14 | 17 | | 180 |
| +10 mM L-lactate | 14 | 19 | 0.0002 | 180 |
| +10 mM pyruvate | 14 | 21 | <0.0001 | 180 |
| ***ire-1(v33)*** | 13 | 17 | | 90 |
| +10 mM L-lactate | 15 | 19 | <0.0001 | 90 |
| +10 mM pyruvate | 15 | 19 | <0.0001 | 90 |
| *atfs-1(gk3094)* | 16 | 20 | | 90 |
| +10 mM L-lactate | 20 | 26 | <0.0001 | 90 |
| +10 mM pyruvate | 16 | 24 | 0,0012 | 90 |
| **HT115-EV** | 15 | 25 | | 90 |
| +10 mM L-lactate | 17 | 23 | 0.0018 | 90 |
| +10 mM pyruvate | 20 | 28 | <0.0001 | 90 |
| ***ldh-1* RNAi** | 17 | 23 | | 90 |
| +10 mM L-lactate | 17 | 23 | n.s. 0.9391 | 90 |
| +10 mM pyruvate | 19 | 25 | 0.0001 | 90 |
| ***slcf-1* RNAi** | 21 | 32 | | 90 |
| +10 mM L-lactate | 23 | 28 | 0.0018 | 90 |
| +10 mM pyruvate | 23 | 30 | <0.0001 | 90 |

**Supplementary Table I**: *C. elegans* lifespans statistics ***(Related to Figures 4 and 6, Supplementary Figure 3 and 6)***
